# Releasing RNA from formalin: novel 20^th^ century rotavirus alphagastroenteritidis strains identified in Australian microbat voucher specimens

**DOI:** 10.1101/2025.09.07.674729

**Authors:** Ashleigh F Porter, Clare E Holleley, Celeste Donato, Erin E Hahn, Ina Smith, Tonya Haff, Christopher A Wilson, Marina R Alexander

## Abstract

Archival specimens held in biorepositories (e.g. natural history collections) offer rare temporal snapshots of global biodiversity. These collections not only preserve species morphology and aspects of ecology, but increasingly provide access to historical molecular data, including insights into wildlife disease. As several pandemics have originated from animal viruses spilling over into the human population (i.e., SARS-CoV-2/COVID-19, 2009 H1N1 influenza, and HIV/AIDS), characterising the diversity of viruses circulating in wildlife populations is essential for proactive pandemic preparedness. Yet, current surveillance remains biased toward contemporary viruses of economic importance. One solution to bridging spatiotemporal gaps in wildlife virus knowledge is retrospective screening of vouchered wildlife specimens. However, such efforts have been hindered by formalin fixation of specimens, which degrades and cross-links nucleic acids. Here we demonstrate that formalin-fixed vouchered wildlife specimens retain both host and viral RNA fragments after being stored for up to sixty years. We recovered fragments of divergent strains of *Rotavirus alphagastroenteritidis* from two Australian microbat species; *Nyctophilus geoffroyi* (lesser long-eared bat) and *Rhinolophus megaphyllus* (smaller horseshoe bat), representing the first characterisation of *Rotavirus alphagastroenteritidis* in Australian bats, and the oldest identification of the virus to date. Concurrently, we sequenced endogenous host RNA, providing a proof-of-concept for dual host-virus transcript recovery from vouchered specimens. This study highlights the role biorepositories can play in reconstructing historical viral landscapes and enabling spatiotemporal host-virus insight to advance both biodiversity science and global pandemic preparedness.

## Introduction

Emerging infectious diseases (EIDs), including zoonotic viruses, cause a substantial burden for human and animal health, with approximately one novel or reemerging threat detected annually (Bloom, et al. 2017). Climate and land use changes (e.g., habitat destruction), resource scarcity (e.g., decreased food availability), and increased interactions between humans, wildlife and domesticated animals have expanded the human-wildlife and wildlife-domestic animal interfaces, and are fuelling the risk of zoonotic viral spillover (Woolhouse and Gaunt 2007; Bloom, et al. 2017; Plowright, et al. 2024).

To date, viral discovery and characterisation efforts have been skewed toward pathogens that affect human health or agricultural productivity. This bias is perpetuated in public genetic sequence repositories, reducing the effectiveness of viral discovery tools that rely upon this data and hindering identification of novel viruses across the broader viral ecosphere (Krishnamurthy and Wang 2017).

Furthermore, the vast majority of wildlife viruses with zoonotic potential remain uncharacterised (Carroll, et al. 2018) and are investigated only once there is sustained circulation in the human population (Vora, et al. 2022). Practical pandemic preparedness requires broad-scale, unbiased, proactive surveillance to advance viral discovery in wildlife (Plowright, et al. 2024).

Natural history collections may offer a powerful, yet underutilised, resource for retrospective disease surveillance (Colella, et al. 2021). Museum and biobank specimens provide unparalleled temporal and taxonomic coverage that has long been used to describe new eukaryotic species and monitor long-term population dynamics and ecological response to environmental change (Suarez and Tsutsui 2004; Cook, et al. 2014; Funk 2018; Meineke, et al. 2019; Nachman, et al. 2023). Notably, most whole specimens collected prior to the 1980s were preserved in formalin, a chemical that crosslinks cellular molecules and was historically thought to prohibit molecular analysis (Simmons 2014).

Recent biochemical and sequencing innovations have overturned the assumption that molecular data from formalin-fixed specimens is inaccessible (Holleley and Hahn 2025). Studies now demonstrate that DNA can be recovered from such material, enabling genomic, phylogenetic, and epigenomic analysis (Hykin, et al. 2015; Ruane and Austin 2017; Hahn, et al. 2020; Totoiu, et al. 2020; Straube, et al. 2021; Hahn, Alexander, et al. 2024; Hahn, Stiller, et al. 2024; Gieraths, et al. 2025; Holleley and Hahn 2025). These advances open new avenues not only for studying host biology but also to investigate the diversity and evolution of wildlife viruses within a single host (e.g., the “virome”) and across populations, through space and time.

Proposed actions to integrate natural history collections and microbiological research to enhance pandemic preparedness (Thompson Cody, et al. 2021) have gained traction following the successful identification of several pathogens in archived specimens, including Sin Nombre virus, monkeypox virus, *Yersinia pestis*, *Mycobacterium tuberculosis, Pseudogymnoascus destructans,* and *Cichlidogyrus spp.* (Mills, et al. 1998; Rothschild, et al. 2001; Yates, et al. 2002; de Oliveira and Franco 2005; Van Steenberge, et al. 2015; Campana, et al. 2017; Tiee, et al. 2018).

Here, we applied museum-tailored RNA extraction methods and metatranscriptomic sequencing to a set of formalin- and ethanol-preserved Australian microbat specimens from the Australian National Wildlife Collection (ANWC), including *Nyctophilus geoffroyi, Rhinolophus megaphyllus, Ozimops ridei, Scotorepens greyii*, *Hipposideros diadema* and *Mops jobensis*. We successfully recovered both host RNA and reads of viral origin, which allowed us to reconstruct fragments of *Rotavirus alphagastroenteritidis* (RVA) from two specimens, representing divergent strains.

These strains are the first RVA detections in Australian bats and include one of the oldest RVA detections worldwide. RVA is a leading cause of acute gastroenteritis, which carries a serious mortality risk in young humans (>120,000 paediatric deaths per annum), agricultural livestock and wild-living animals (Adams and Kraft 1963; Rodger, et al. 1975; Mebus 1976; Tate, et al. 2012; Lanata, et al. 2013; Aboyans and Collaborators 2015). Our findings demonstrate that formalin-fixed specimens can retain both viral and host RNA for over sixty years, highlighting the potential of museum collections to contribute to long-term viral surveillance, expand our knowledge of viral biodiversity and inform future studies in host-pathogen evolution, biodiversity, and global health.

## Results

### Archival RNA of host and viral origin is retrievable from museum specimens

We successfully extracted RNA from seven vouchered microbat specimens preserved in either formalin or ethanol (Table 1). Despite chemical fixation and specimen ages exceeding 60 years, tissue integrity was sufficient for specimen morphology and dissection (Figure 1a,b) and to enable RNA recovery (Figure 1e,f). RNA yields ranged from 2.9 ng/μL to 48.4 ng/μL (Figure 1e; Table 1) and all samples produced libraries with suitable read counts and lengths for downstream analysis (Figure 1f; Supplementary Table 1). Post-trimming quality scores exceeded Q34, and no sequences were flagged as poor quality by FASTQC.

**Figure 1.**
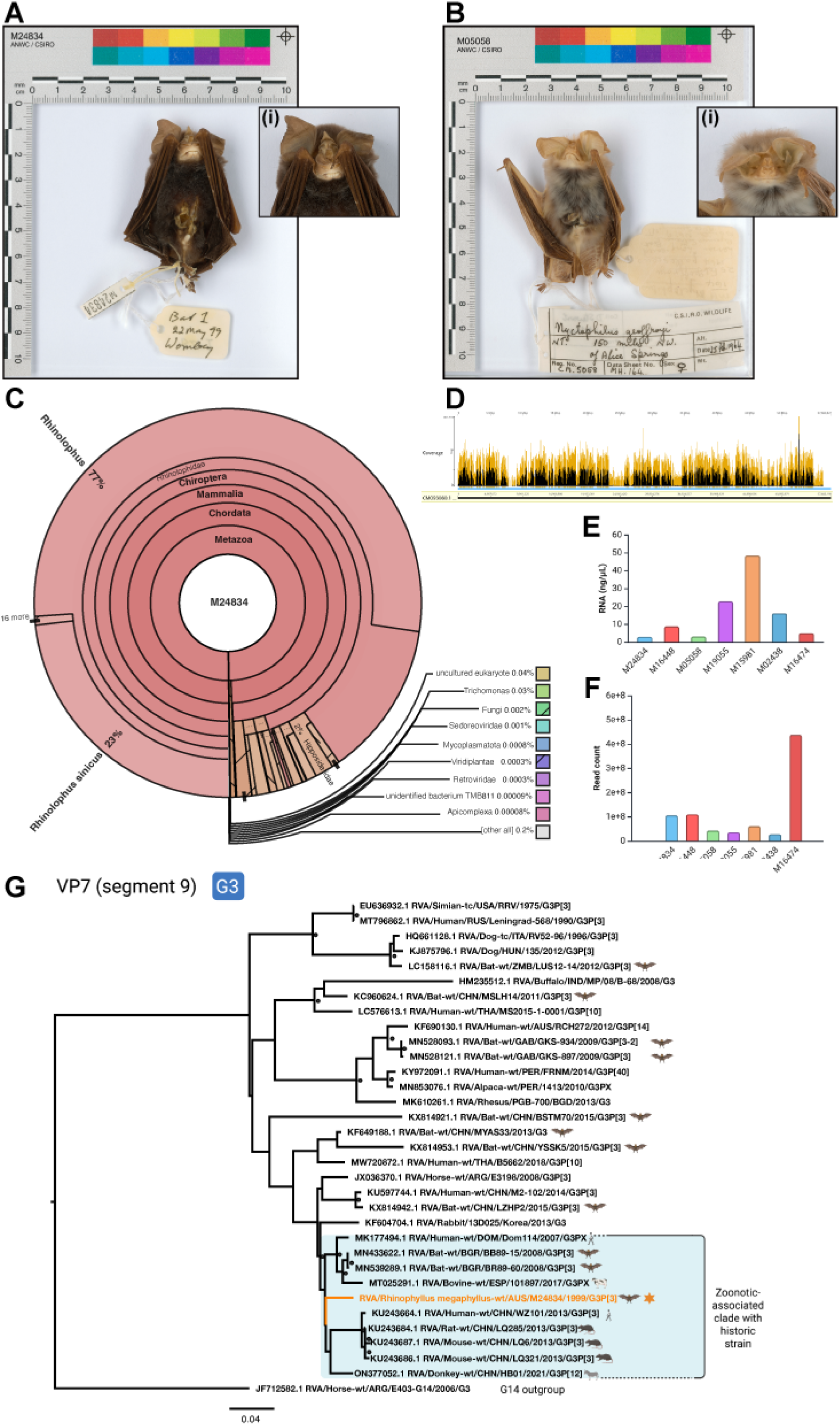
Specimen condition, RNA-seq output, and phylogenetic placement of novel historical strain. (A) Anterior view of specimen M24834 (*Rhinolophus megaphyllus*), an adult male smaller horseshoe bat with a fleshy horseshoe-shaped nose leaf and grey-brown fur. Body mass at collection was 8.8g. The nose leaf can be observed in close-up (A[i]). (B) Anterior view of specimen M05058 (*Nyctophilus geoffroyi*), an adult female lesser long-eared bat with pale grey fur on the back and white fur on the underbelly. Forearm length at collection was 30.4mm. The facial features can be observed in close-up (B[i]). (C) Taxonomic origin of reads inferred using the NCMI nt database visualised for specimen M24834. (D) Visualisation of trimmed reads from specimen M24834 mapped to the *Rhinolophus megaphyllus* genome (CM093080.1, chromosome 18). (E) Total RNA yield (ng/μL) recovered from liver tissue per specimen. (F) Total RNA-seq read count recovered per specimen. (G) Maximum likelihood phylogeny of VP7 (segment 9) sequences from genotype G3 RVA strains (n=41). The novel historical strain RVA/Rhinolophus megaphyllus- wt/AUS/M24834/1999/G3/P[3] is highlighted in orange with a star. Tip icons indicate host origin (e.g., bat, human, mouse) and blue boxes denote the clade containing the historical strain. Bootstrap values > 80% are shown as filled circles and the scale bar represents nucleotide substitutions per site. The phylogeny is mid-point rooted using a G14 outgroup.

**Table 1.** Details of Australian microbat specimens screened for viral RNA. We screened seven specimens from the Australian National Wildlife Collection (ANWC) for viral RNA using liver tissue samples. For each specimen, we report the ANWC specimen registration number, species name (scientific and common), collection location (Australian state), collection date, preservation method, library preparation approach, and total RNA yield (ng/μL) as measured by Agilent 2100 Bioanalyzer. Bolded specimens yielded viral hits discussed in this study. NT, Northern Territory; QLD, Queensland.

| Registration number | Species name | Location | Collection date | Preservation method | Library preparation methodology | RNA (yield ng/μL) |
| --- | --- | --- | --- | --- | --- | --- |
| **ANWC M05058** | ***Nyctophilus geoffroyi***<br>(lesser long eared bat) | NT | 25-2-1964 | Formalin-fixed and kept at room temperature | Total nucleic acid library preparation and targeted enrichment using Twist probes | 3.1 |
| **ANWC M24834** | ***Rhinolophus megaphyllus*** (smaller horseshoe bat) | QLD | 22-5-1999 | Ethanol-fixed and frozen at -20°C after collection | Illumina TruSeqStranded mRNA prep | 2.9 |
| ANWC M16448 | *Ozimops ridei* or *Mormopterus loriae* (eastern free tailed bat) | QLD | 23-2-1992 | Ethanol-fixed and frozen at -20°C after collection | Illumina TruSeqStranded mRNA prep | 8.8 |
| ANWC M15981 | *Scotorepens greyii* (Little broad-nosed bat) | QLD | 25-2-1992 | Formalin-fixed and kept at room temperature | Total nucleic acid library preparation and targeted enrichment using Twist probes | 48.4 |
| ANWC M16474 | *Hipposideros diadema* (Diadem leaf-nosed bat) | QLD | 7-2-1966 | Formalin-fixed and kept at room temperature | Total nucleic acid library preparation and targeted enrichment using Twist probes | 4.9 |
| ANWC M02438 | *Nyctophilus geoffroyi* (lesser long eared bat) | NT | 26-4-1968 | Formalin-fixed and kept at room temperature | Total nucleic acid library preparation and targeted enrichment using Twist probes | 16.1 |
| ANWC M19055 | *Mops jobensis* (Northern freetail bat) | NT | 30-6-1974 | Formalin-fixed and kept at room temperature | Total nucleic acid library preparation and targeted enrichment using Twist probes | 22.8 |

Across libraries, a high proportion of reads (mean 87%) were repetitive or consisted of short k-mer dimers (<30bp) of unknown origin (sequencing duplication, microsatellites, promoter regions, repetitive elements), indicating that only a subset represented identifiable endogenous host or microbial RNA (Supplementary Figure 1, Supplementary Table 2). Nevertheless, taxonomic profiling revealed in two specimens, ANWC M24834 and M16448 (all specimens from ANWC unless otherwise noted), that most assignable reads originated from the host (Figure 1c, Supplementary Figure 2), with smaller fractions attributed to other eukaryotes, fungi, and viruses (Figure 1c; Supplementary Figure 2). In specimen M24834, viral reads were attributed to the families Sedoreoviridae (containing the *Rotavirus* genus) and Retroviridae. Three specimens, M16474, M19055, and M15981, had the majority of reads assigned to order Mammalia, with a small proportion assigned to Chiroptera, while specimen M02438 and M15981 had almost half of reads unassigned (Supplementary Figure 2) and M05058 had 17% of reads map to Mammalia, 42% map to viral order Reovirales (containing Sedoreoviridae), and 26% of reads unassigned.

We mapped reads from specimen M24834 (*Rhinolophus megaphyllus*) to the *R. affinis* genome, the closest available annotated genome, and found that 62.58% of reads aligned uniquely across all 32 chromosomes (Figure 1d, Supplementary Table 3). For other species lacking reference genomes, we mapped to partial host gene sequences, with lower alignment scoring compared to whole annotated genome alignment. For example, 0.03% of reads from specimen M16448 mapped to a 759 bp NADH gene fragment from *Ozimops ridei* (Supplementary Table 4). In three specimens (M16474, M19055, and M15981), we could not map any reads to a host gene.

We constructed virus-like contigs (e.g., *de novo* sequences with significant homology to viral genes) from sequencing libraries from three specimens (M24834, M05058, and M16448). In M24834 and M05058, most viral contigs matched rotavirus alphagastroenteritidis (RVA), while additional contigs from M24834 resembled bat gammaherpesvirus and bat faecal-associated retrovirus. In M16448, a subset of contigs showed similarity to *Rhinolophus ferrumequinum* gammaretrovirus. From M24834 and M05058, we assembled 16 candidate RVA contigs, ranging from 129 to 2,508 bp in length (Table 2). The relative abundance of RVA-like reads in these libraries ranged from 0.00001% to 6.1% (Supplementary Table 2).

**Table 2.** Summary of viral genome segments recovered from two formalin-preserved bat museum specimens. For each specimen, we list the RVA strain name, genomic segment with corresponding protein, assigned genotype, GC content (%GC), and contiguous sequence length (bp). Genotypes are based on established classification criteria for RVA. Complete coding sequences (cds) are indicated where recovered. Sequences with gaps – represented by stretches of ambiguous bases (‘Ns’) – are noted with a superscript hash (#). Genotypes marked with an asterisk (*) could not be confidently assigned. VP – viral protein; NSP – non-structural protein

| Strain | Segment and protein | Genotype | %GC | Sequence Length (bp) |
| --- | --- | --- | --- | --- |
| RVA/Nyctophilus geoffroyi-wt/AUS/M05058/1964/G3PX | Segment 2 (VP2), partial | C2* | 39.6% | 321 |
|  | Segment 4 (VP4), partial | P[3]* | 37.2% | 234 |
|  | Segment 6 (VP6), partial | I3 | 42.3% | 657 |
|  | Segment 9 (VP7), partial | G3 | 37.8% | 870<sup>#</sup> |
|  | Segment 8 (NSP2),<br>partial | n/a | 45.0% | 129 |
|  | Segment 11 (NSP5),<br>partial | H6 | 39.6% | 492 |
| RVA/Rhinolophus<br>megaphyllus-wt/AUS/M24384/1999/G3P[3] | Segment 1 (VP1), partial | R3 | 33.3% | 3,267<sup>#</sup> |
|  | Segment 2 (VP2), cds | C3 | 33.6% | 2,664 |
|  | Segment 3 (VP3), cds | M3 | 30.9% | 2,508 |
|  | Segment 4 (VP4), partial | P[3] | 37.1% | 1,896<sup>#</sup> |
|  | Segment 5 (NSP1),<br>partial | A9 | 33.3% | 1,395 |
|  | Segment 6 (VP6), partial | I3 | 33.5% | 564 |
|  | Segment 7 (NSP3),<br>partial | T3 | 34.1% | 954 |
|  | Segment 8 (NSP2),<br>partial | N3 | 33.5% | 477 |
|  | Segment 9 (VP7), cds | G3 | 35.2% | 981 |
|  | Segment 11 (NSP5),<br>partial | H6 | 36.3% | 408 |

### Assembly of historical RVA genomes reveals specimen and method-specific variation

Using a metatranscriptomic approach, we recovered historical viral RNA from two chemically preserved specimens (M24834 and M05058) collected in 1999 and 1964, respectively. Both yielded sequences from *Rotavirus alphagastroenteritidis* (RVA), a double-stranded RNA virus of the family Sedoreoviridae, comprising 11 genomic segments that encode structural (VP1 – VP6) and non-structural proteins (NSP1 – NSP6) (Estes, et al. 2007). We detected RVA using two complementary methods in different specimens; total RNA-seq (M24834) and targeted enrichment (M05058) using the Twist Comprehensive Viral Research Panel, supplemented with custom probes targeting Gammaretrovirus (see Supplementary Methods, Supplementary Table 5).

From specimen M24834 (*R. megaphyllus*), we recovered three coding sequences (cds) (VP2, VP3, and VP7) and partial sequences for VP1, VP4, VP6, NSP1, NSP2, NSP3, and NSP5 (Table 2). The VP1 and VP4 segments were assembled from multiple contigs, with ambiguous bases (‘Ns’) inserted to span unsupported gaps. In specimen M05058 (*N. geoffroyi*), we reconstructed partial sequences for VP2, VP4, VP6, NSP2, NSP5, and VP7. All but VP7 were recovered as single contigs while VP7 was assembled from two contigs bridged with 257 ambiguous bases (Table 2).

We note that the RVA sequences obtained from specimen M05058 were generally shorter and more fragmented than those from specimen M24834. This discrepancy may reflect differences in viral load, specimen preservation quality, or biases introduced by the probe set, which was not specifically optimised for Sedoreoviridae. Despite inclusion of retroviral probes in our enrichment panel, we did not detect Retroviridae sequences in any of the five probe-enriched libraries, only a small proportion of Retroviridae reads that did not assemble into large contigs were detected in the total RNA libraries.

### Rotavirus A genotypes reveal reassortment and potential host connections

RVA exhibits high genetic diversity driven by two evolutionary mechanisms: genetic drift and genetic shift. Genetic drift arises through the accumulation of point mutations introduced by the virus’s error-prone RNA-dependent RNA polymerase (Ramig 1997), although some mutations are retained through positive selection. In contrast, genetic shift occurs through reassortment of two or more distinct RVA strains during co-infection of a host cell, allowing gene segment exchange; a key mechanism driving viral diversification and interspecies transmission (Matthijnssens, Ciarlet, Heiman, et al. 2008; Martella, et al. 2010; Matthijnssens, De Grazia, et al. 2011).

To characterise viral diversity, RVA is classified using a dual genotyping system based on the outer capsid proteins VP7 and VP4 genes. These define the G- genotype (VP7 is g<u>lycosylated)</u> and P-genotype (VP4 is p<u>rotease</u> sensitive). A more comprehensive classification known as <u>genotype constellation</u> assigns genotypes to all 11 genome segments using a standard nomenclature: Gx – P[x] – Ix – Rx – Cx – Mx – Ax – Nx – Tx – Ex – Hx, representing VP7 – VP4 – VP6 – VP1 – VP2 – VP3 – NSP1 – NSP2 – NSP3 – NSP4 – NSP5 (Matthijnssens, Ciarlet, et al. 2011).

We reconstructed partial or complete genotype constellations for both museum-derived strains. Historical strain RVA/Rhinolophus megaphyllus- wt/AUS/M24384/1999/G3P[3] demonstrated the constellation G3 – P[3] – I3 – R3 – C3 – M3 – A9 – N3 – T3 – Ex – H6 (Matthijnssens, Ciarlet, et al. 2011), which we confirmed using the Rotavirus A Genotyping Tool, v1 (rivm.nl/mpf/typingtool/rotavirusa). Four segments (VP4, NSP2, NSP4, and NSP5) fell below the 500 bp threshold for confident classification and thus are considered tentative (Matthijnssens, Ciarlet, et al. 2011). This Australian genotype constellation is identical to bat-derived strains from Bulgaria (e.g., RVA/Bat-wt/BGR/BB89- 15/2008/G3P[3]; RVA/Bat-wt/BGR/BR89-60/2008/G3P[3]).

In contrast, the historical strain RVA/Nyctophilus geoffroyi- wt/AUS/M05058/1964/G3PX strain exhibited a constellation of: G3 – P[3]* – I3* – Rx – C2* – Mx – Ax – Nx – Tx – Ex – H6, with asterisks indicating genotypes (VP4, VP6, NSP2) inferred from short sequences (Matthijnssens, Ciarlet, et al. 2011). As such, the VP4 genotype remains unassigned. Segment 10 (NSP4) was not recovered, and three segments (VP2, VP4, NSP5) were < 500 bp and considered incomplete. This partial genotype constellation resembles early zoonotic strains reported in humans, including RVA/Human/ISR/Ro1845/1985/G3P[3] and RVA/Human/USA/HCR3A/1984/G3P[3], which were originally thought to be of canine or feline origin (Aboudy, et al. 1988; Nakagomi, et al. 1990; Li, et al. 1993; Nakagomi and Nakagomi 2000; Tsugawa and Hoshino 2008).

We visualised recovered segments for both strains relative to a canonical segmented RVA genome (Figure 2a) and identified their closest relatives using megablast nucleotide identity (Supplementary Table 6).

**Figure 2.**
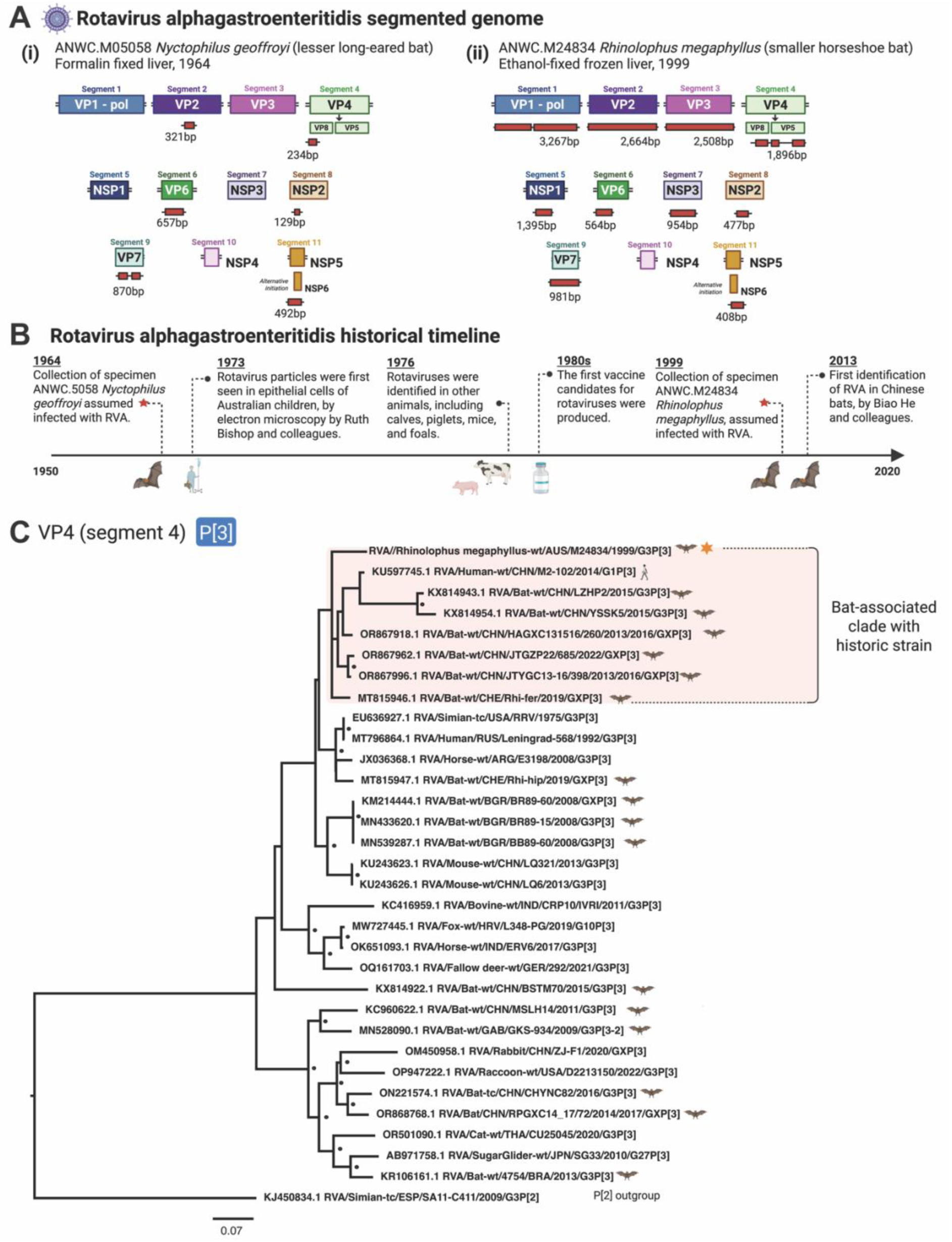
Genome characterisation of two novel bat RVA strains, historic timeline of RVA, and phylogenetic placement of strain RVA/Rhinolophus megaphyllus-wt/AUS/M24384/1999/G3P[3] within VP4 phylogeny. (A) Partial assemblies of RVA strains recovered from two archived bat specimens: [i] *Nyctophilus geoffroyi* (M05058, 1964) and [ii]*Rhinolophus megaphyllus* (M24834, 1999). Segments of the canonical RVA genome are shown as coloured boxes and annotated with segment and protein names. Recovered contigs are shown as red boxes aligned by segment, scaled to approximate length and placement. (B) A historical timeline displaying major events in RVA discovery throughout the last 70 years. Left to right: 1964; the collection of specimen M05058, *Nyctophilus geoffroyi*, assumed to be infected with RVA, 1973; Ruth Bishop and colleagues discover the particles that will be termed “rotavirus” under the electron microscope, seen in epithelial cells obtained from Australian patients, 1976; Rotaviruses are also identified in other mammals, including cows, pigs, mice and horses, 1980s; vaccine development for rotavirus begins, 1999; the collection of specimen M24834, *Rhinolophus megaphyllus*, assumed to be infected with RVA, 2013; first modern detection of RVA in bats by Biao He and colleagues in China. (C) Phylogenetic tree of RVA segment 4/VP4, containing a 450bp section from the partial sequence (three contigs joined by empty strings) from strain RVA/Rhinolophus megaphyllus- wt/AUS/M24384/1999/G3P[3], along with other partial or whole coding sequences of RVA VP4. genotype P[3] (n=32). The novel historical strain has the respective branch highlighted in orange and an orange star next to the tip name. Bat-associated strains have a bat icon next to their tip name. Strains within the same clade as the novel historical strain have their associated animal host icon next to their tip name. To visualise the grouping of bat strains and the historical strain, the clade has been highlighted by a red box. Bootstrap replicates > 80% are visualised as filled-in circles at nodes. The scale bar represents nucleotide substitutions per site. For clarity, the phylogeny is rooted at the P[2] outgroup.

### 20th century bat RVA strains point to long-standing zoonotic potential

Rotaviruses infect many species and RVA circulates widely across livestock, companion animals, and wildlife, forming a diverse reservoir with potential for interspecies transmission (Matthijnssens, Ciarlet, Heiman, et al. 2008; Ghosh and Kobayashi 2014; Lagan, et al. 2023). In humans, RVA lineages are thought to emerge through repeated zoonotic introductions and reassortment events, as evidenced by sporadic spillovers from bovine and porcine hosts (Chandler-Bostock, et al. 2014; Desselberger 2014; Phan, et al. 2016; Zeller, et al. 2016; Sawant, et al. 2020).

Bats are frequent hosts of RVA, harbouring a wide diversity of genotypes. Some of these genotypes appear bat-specific, while others are shared with alternate mammalian hosts including humans, suggesting a history of cross-species transmission (Esona, et al. 2010; He, Yang, Yang, Zhang, Feng, Zhou, Xie, Feng, Bao and Guo 2013; Asano, et al. 2016; Sasaki, et al. 2016; Yinda, et al. 2016; He, et al. 2017; Simsek, et al. 2021). Bat-associated RVA strains have been identified around the world in Saudi Arabia, Zambia, Brazil, China, Cameroon, Kenya, Bangladesh, and across Europe in multiple bat families including Pteropodidae, Rhinolophidae, Hipposideridae, Pyllostomidae, Vespertilionidae, Emballonuridae, and Miniopteridae (He, Yang, Yang, Zhang, Feng, Zhou, Xie, Feng, Bao, Guo, et al. 2013; Dacheux, et al. 2014; Xia, et al. 2014; Asano, et al. 2016; Sasaki, et al. 2016; Yinda, et al. 2016; He, et al. 2017; Waruhiu, et al. 2017; Mishra, et al. 2019; Islam, et al. 2020; Simsek, et al. 2021).

Like zoonotic influenza viruses or other rotaviruses, bat RVA strains can become “humanised” through reassortment with co-circulating human strains (Matthijnssens, et al. 2010; Li, et al. 2016). For example, RVA/Human-wt/CHN/WZ101/2013/G3P[3] closely clustered with rodent-associated strains from the same geographic region, providing evidence of zoonotic spillover. The historical bat strains recovered in our study also cluster phylogenetically with animal-derived RVA strains, further supporting their animal origin.

In Australia, a putatively zoonotic strain RVA/Human-wt/AUS/RCH272/2012/G3P[14] was isolated from a 12-year-old child with gastroenteritis in Victoria in 2012. The individual lived in a household with domesticated cats and dogs, in close proximity to a grey-headed flying fox (*Pteropus poliocephalus)* colony. This strain exhibited a mosaic genome, with segments resembling international bat strains and lineages of bovine, canine, feline, human origins. However, no Australian bat RVA strains were available for comparison at the time (Figure 2b) (Donato, et al. 2014). While the novel strains discussed here do not cluster with RVA/Human-wt/AUS/RCH272/2012/G3P[14], knowledge of the unsampled diversity of RVA strains within Australian bat populations is critical for future genomic case definitions (particularly zoonoses) and public health interventions.

### Expanding the known diversity of RVA in Australian bats

Bats are among the most species-rich orders of mammals, with over 1,400 species across 21 families (Burgin, et al. 2018). Australian bats span both suborders of Chiroptera: Yangochiroptera (primarily microbats) and Yinpterochiroptera (including Pteropodidae, or megabats, and five microbat families). Bats host a broad array of viruses, many still undescribed (Tan, et al. 2021), and several with zoonotic potential, including henipaviruses, lyssaviruses, filoviruses, and coronaviruses. As a result, bats have become a focal group for viral surveillance and virome characterisation (Wu, et al. 2016; Van Brussel and Holmes 2022; Wallau, et al. 2023).

RVA is frequently detected in bat faecal samples, with over 20 bat-associated lineages now described globally (Esona, et al. 2010; He, Yang, Yang, Zhang, Feng, Zhou, Xie, Feng, Bao and Guo 2013; Xia, et al. 2014; Asano, et al. 2016; Sasaki, et al. 2016; Yinda, et al. 2016; He, et al. 2017; Waruhiu, et al. 2017; Mishra, et al. 2019; Simsek, et al. 2021). The sequences presented here represent the first RVA strains identified and characterised from Australian bats. Although previous studies speculated that Australian flying foxes (Pteropodidae) may act as RVA reservoirs (Donato, et al. 2014), our data establish the first confirmed Australian bat-associated RVA strains from microbats. Given the high bat diversity in Australia, it is likely that bat-associated RVA diversity in this region is far greater than currently recognised.

These findings establish an initial baseline for RVA diversity in Australian bats and highlight the untapped potential of museum specimens for revealing previously undocumented viral lineages.

### Phylogenetic relationships reveal global links and reassortment in historical bat RVA

To contextualise the evolutionary history of the recovered strain RVA/Rhinolophus megaphyllus-wt/AUS/M24384/1999/G3P[3], we conducted maximum likelihood phylogenetic analyses for each segment using representative RVA sequences (models in Supplementary Table 7). This strain clustered with zoonosis-associated clades in the VP1, VP6, VP7, NSP1, and NSP2 trees (Figure 1g, Supplementary Figures 3, 6-8), suggesting broad host plasticity and potential for interspecies transmission. In contrast, segments VP4 and NSP3 clustered exclusively with other bat-associated strains (Figure 2c, Supplementary Figure 9), indicating more host-restricted evolutionary trajectories for these segments.

We identified a genotype constellation shared between the novel strain RVA/AUS/Rhinolophus megaphyllus-wt/M24834/1999/G3/P[3] and a subset of published sequences, which we refer to here as the “Au/zoo” constellation (Table 3). This constellation has been isolated from a broad array of hosts, including bats (*Rhinolophus* spp.), rodents (e.g., reed voles), horses, and humans, and has been detected multiple times over more than a decade (1999 - 2015) in China, Bulgaria, Argentina and now Australia. Its recurrence in multiple hosts and geographic contexts suggests either long-term circulation in reservoir populations or repeated emergence through reassortment.

**Table 3.**
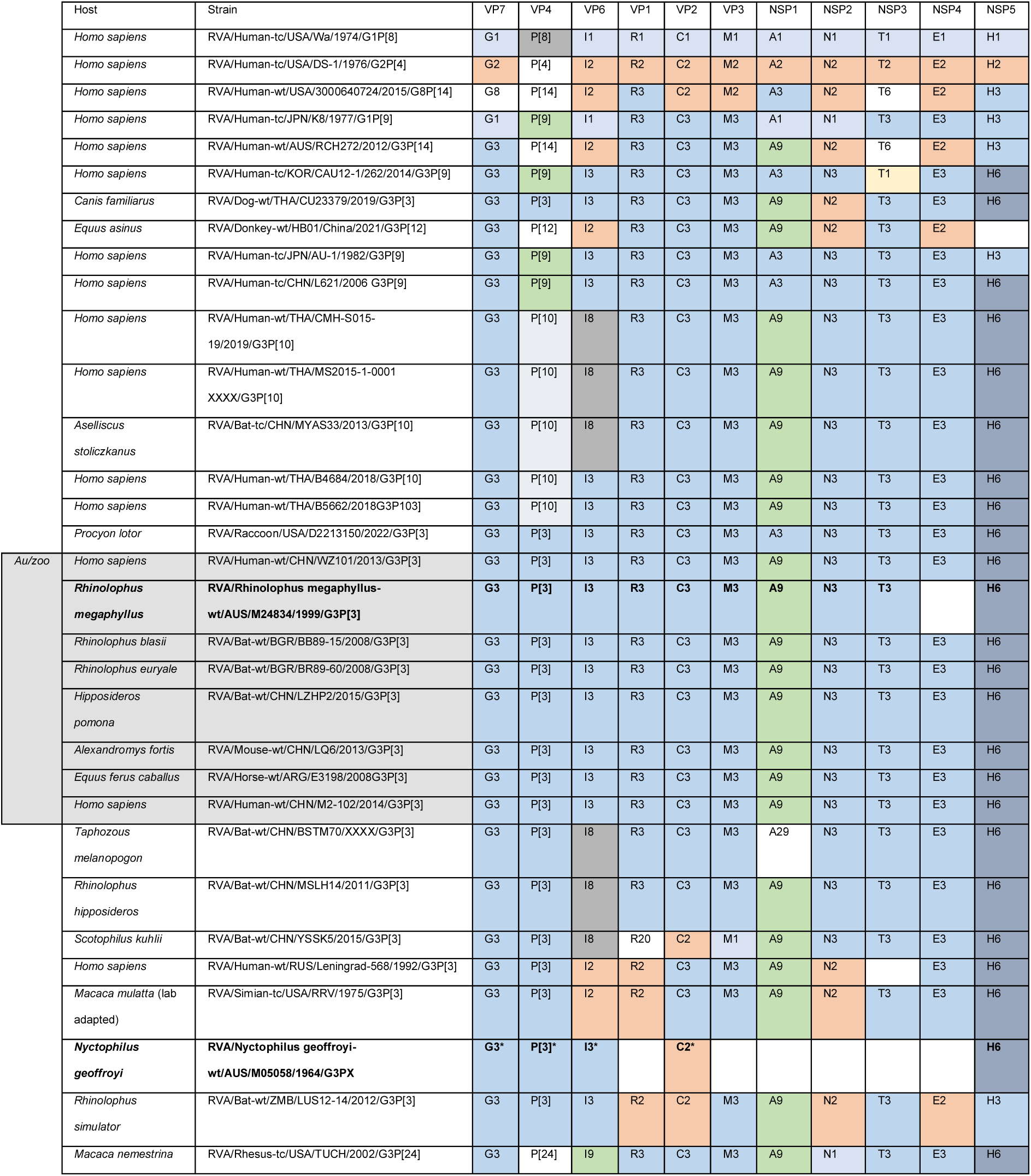
Genotype constellations of novel and related RVA strains across hosts. We summarise the genotype constellations for RVA/Nyctophilus geoffroyi- wt/AUS/M05058/1964/G3PX, RVA/Rhinolophus megaphyllus- wt/AUS/M24384/1999/G3P[3] and other well known RVA strains, and strains that share genotypes. For each strain, we indicate the host species (scientific name), strain designation, and assigned genotype for all 11 RVA genome segments (VP7, VP4, VP6, VP1, VP2, VP3, NSP1, NSP2, NSP3, NSP4, and NSP5). Shared genotypes are colour-coded to visualise patterns of genomic similarity and reassortment. Strains included in the “Au/zoo” genotype constellation are grouped within the grey-shaded box. Genotypes marked with an asterisk (*) are uncertain due to contig length or sequence identity thresholds.

|  | Host | Strain | VP7 | VP4 | VP6 | VP1 | VP2 | VP3 | NSP1 | NSP2 | NSP3 | NSP4 | NSP5 |
| --- | --- | --- | --- | --- | --- | --- | --- | --- | --- | --- | --- | --- | --- |
|  | *Homo sapiens* | RVA/Human-tc/USA/Wa/1974/G1P[8] | G1 | P[8] | I1 | R1 | C1 | M1 | A1 | N1 | T1 | E1 | H1 |
|  | *Homo sapiens* | RVA/Human-tc/USA/DS-1/1976/G2P[4] | G2 | P[4] | I2 | R2 | C2 | M2 | A2 | N2 | T2 | E2 | H2 |
|  | *Homo sapiens* | RVA/Human-wt/USA/3000640724/2015/G8P[14] | G8 | P[14] | I2 | R3 | C2 | M2 | A3 | N2 | T6 | E2 | H3 |
|  | *Homo sapiens* | RVA/Human-tc/JPN/K8/1977/G1P[9] | G1 | P[9] | I1 | R3 | C3 | M3 | A1 | N1 | T3 | E3 | H3 |
|  | *Homo sapiens* | RVA/Human-wt/AUS/RCH272/2012/G3P[14] | G3 | P[14] | I2 | R3 | C3 | M3 | A9 | N2 | T6 | E2 | H3 |
|  | *Homo sapiens* | RVA/Human-tc/KOR/CAU12-1/262/2014/G3P[9] | G3 | P[9] | I3 | R3 | C3 | M3 | A3 | N3 | T1 | E3 | H6 |
|  | *Canis familiaris* | RVA/Dog-wt/THA/CU23379/2019/G3P[3] | G3 | P[3] | I3 | R3 | C3 | M3 | A9 | N2 | T3 | E3 | H6 |
|  | *Equus asinus* | RVA/Donkey-wt/HB01/China/2021/G3P[12] | G3 | P[12] | I2 | R3 | C3 | M3 | A9 | N2 | T3 | E2 |  |
|  | *Homo sapiens* | RVA/Human-tc/JPN/AU-1/1982/G3P[9] | G3 | P[9] | I3 | R3 | C3 | M3 | A3 | N3 | T3 | E3 | H3 |
|  | *Homo sapiens* | RVA/Human-tc/CHN/L621/2006 G3P[9] | G3 | P[9] | I3 | R3 | C3 | M3 | A3 | N3 | T3 | E3 | H6 |
|  | *Homo sapiens* | RVA/Human-wt/THA/CMH-S015-19/2019/G3P[10] | G3 | P[10] | I8 | R3 | C3 | M3 | A9 | N3 | T3 | E3 | H6 |
|  | *Homo sapiens* | RVA/Human-wt/THA/MS2015-1-0001 XXXX/G3P[10] | G3 | P[10] | I8 | R3 | C3 | M3 | A9 | N3 | T3 | E3 | H6 |
|  | *Aselliscus stoliczkanus* | RVA/Bat-tc/CHN/MYAS33/2013/G3P[10] | G3 | P[10] | I8 | R3 | C3 | M3 | A9 | N3 | T3 | E3 | H6 |
|  | *Homo sapiens* | RVA/Human-wt/THA/B4684/2018/G3P[10] | G3 | P[10] | I3 | R3 | C3 | M3 | A9 | N3 | T3 | E3 | H6 |
|  | *Homo sapiens* | RVA/Human-wt/THA/B5662/2018G3P103] | G3 | P[10] | I3 | R3 | C3 | M3 | A9 | N3 | T3 | E3 | H6 |
|  | *Procyon lotor* | RVA/Raccoon/USA/D2213150/2022/G3P[3] | G3 | P[3] | I3 | R3 | C3 | M3 | A3 | N3 | T3 | E3 | H6 |
| Au/zoo | *Homo sapiens* | RVA/Human-wt/CHN/WZ101/2013/G3P[3] | G3 | P[3] | I3 | R3 | C3 | M3 | A9 | N3 | T3 | E3 | H6 |
|  | *Rhinolophus megaphyllus* | RVA/Rhinolophus megaphyllus-wt/AUS/M24834/1999/G3P[3] | G3 | P[3] | I3 | R3 | C3 | M3 | A9 | N3 | T3 |  | H6 |
|  | *Rhinolophus blasii* | RVA/Bat-wt/BGR/BB89-15/2008/G3P[3] | G3 | P[3] | I3 | R3 | C3 | M3 | A9 | N3 | T3 | E3 | H6 |
|  | *Rhinolophus euryale* | RVA/Bat-wt/BGR/BR89-60/2008/G3P[3] | G3 | P[3] | I3 | R3 | C3 | M3 | A9 | N3 | T3 | E3 | H6 |
|  | *Hipposideros pomona* | RVA/Bat-wt/CHN/LZHP2/2015/G3P[3] | G3 | P[3] | I3 | R3 | C3 | M3 | A9 | N3 | T3 | E3 | H6 |
|  | *Alexandromys fortis* | RVA/Mouse-wt/CHN/LQ6/2013/G3P[3] | G3 | P[3] | I3 | R3 | C3 | M3 | A9 | N3 | T3 | E3 | H6 |
|  | *Equus ferus caballus* | RVA/Horse-wt/ARG/E3198/2008G3P[3] | G3 | P[3] | I3 | R3 | C3 | M3 | A9 | N3 | T3 | E3 | H6 |
|  | *Homo sapiens* | RVA/Human-wt/CHN/M2-102/2014/G3P[3] | G3 | P[3] | I3 | R3 | C3 | M3 | A9 | N3 | T3 | E3 | H6 |
|  | *Taphozous melanopogon* | RVA/Bat-wt/CHN/BSTM70/XXXX/G3P[3] | G3 | P[3] | I8 | R3 | C3 | M3 | A29 | N3 | T3 | E3 | H6 |
|  | *Rhinolophus hipposideros* | RVA/Bat-wt/CHN/MSLH14/2011/G3P[3] | G3 | P[3] | I8 | R3 | C3 | M3 | A9 | N3 | T3 | E3 | H6 |
|  | *Scotophilus kuhlii* | RVA/Bat-wt/CHN/YSSK5/2015/G3P[3] | G3 | P[3] | I8 | R20 | C2 | M1 | A9 | N3 | T3 | E3 | H6 |
|  | *Homo sapiens* | RVA/Human-wt/RUS/Leningrad-568/1992/G3P[3] | G3 | P[3] | I2 | R2 | C3 | M3 | A9 | N2 |  | E3 | H6 |
|  | *Macaca mulatta* (lab adapted) | RVA/Simian-tc/USA/RRV/1975/G3P[3] | G3 | P[3] | I2 | R2 | C3 | M3 | A9 | N2 | T3 | E3 | H6 |
|  | *Nyctophilus geoffroyi* | RVA/Nyctophilus geoffroyi-wt/AUS/M05058/1964/G3PX | G3* | P[3]* | I3* |  | C2* |  |  |  |  |  | H6 |
|  | *Rhinolophus simulator* | RVA/Bat-wt/ZMB/LUS12-14/2012/G3P[3] | G3 | P[3] | I3 | R2 | C2 | M3 | A9 | N2 | T3 | E2 | H3 |
|  | *Macaca nemestrina* | RVA/Rhesus-tc/USA/TUCH/2002/G3P[24] | G3 | P[24] | I9 | R3 | C3 | M3 | A9 | N1 | T3 | E3 | H6 |

Phylogenetic comparisons showed that the strain RVA/AUS/Rhinolophus megaphyllus-wt/M24834/1999/G3/P[3] clustered with two human strains, including RVA/Human-wt/CHN/WZ101/2013/G3P[3], a suspected zoonotic strain, and three related rodent strains from the same region (RVA/Rat-wt/CHN/LQ285/2013/G3P[3], RVA/Mouse-wt/CHN/LQ6/2013/G3P[3], and RVA/Mouse- wt/CHN/LQ321/2013/G3P[3]) (Li, et al. 2016). This pattern was consistent across multiple gene trees, including VP7 (Figure 1g), VP6 (Supplementary Figure 7), NSP1 (Supplementary Figure 6), and NSP2 (Supplementary Figure 8). A second human strain, RVA/Human-wt/CHN/M2-102/2014/G3P[3], suspected to have arisen from wildlife reassortment (Dong, et al. 2016), also clustered with the historical strain in the VP4 (Figure 2c), VP1 (Supplementary Figure 3), NSP2, and NSP3 phylogenies.

Two previously reported bat strains RVA/Bat-wt/BGR/BB89-15/2008/G3P[3] and RVA/Bat-wt/BGR/BR89-60/2008/G3P[3], isolated from Bulgarian *Rhinolophus* spp. (Simsek, et al. 2021), also clustered with the Australian strain in VP6 and VP7 phylogenies, strengthening evidence for a long-standing, geographically widespread lineage within *Rhinolophus*-associated RVA. An equine strain with evidence of multiple interspecies transmission and/or reassortment events, RVA/Horse- wt/ARG/E3198/G3P[3], also clustered with the Australian strain in the VP6 and VP3 phylogenies (Miño, et al. 2013).

Due to short length (<300bp) and high divergence of most segments (<80% nucleotide identity), we excluded strain RVA/Nyctophilus geoffroyi- wt/M05058/1964/G3PX from phylogenetic analysis except for VP2 and NSP5 (Matthijnssens, Ciarlet, et al. 2011). Further investigation is required to fully characterise this genotype constellation.

### Historical bat rotaviruses reveal deep evolutionary connections and surveillance gaps

We recovered two distinct partial historical RVA genomes from fragmented RNA in formalin-fixed microbat specimens. To our knowledge, these are the first viral genomes successfully reconstructed from such specimens and the first documented RVA strain from an Australian bat. Remarkably, one historical strain (RVA/Nyctophilus geoffroyi-wt/AUS/M05058/1964/G3PX) predates the discovery of human rotavirus in 1973 (Figure 2b), placing it among the oldest RVA strains characterised to date. Given the absence of prior Australian bat RVA sequences, these genomes may represent endemic strains or ancestral forms of contemporary RVA strains. Both scenarios underscore a significant gap in our understanding of global RVA evolution and geographic spread.

We provide genomic coverage, read abundances (Figure 2a, Supplementary Table 2), genotypes (Table 3), and phylogenetic relationships (Figure 1g, 2c, Supplementary Figures 3-10) for both strains. Viral recovery was not limited by total RNA yield (2.9 ng/μL for M24834 and 3.1 ng/μL for M05058), implicating viral load and specimen preservation quality as more influential factors. The near-complete recovery from M24384 was especially striking, given the specimen’s age and preservation in formalin (Figure 1a). Notably, we recovered substantial host RNA from libraries prepared without enrichment (M24834 and M16448; Figure 1, Supplementary Figure 2), consistent with expectations that probe-based enrichment introduces biases against endogenous transcripts. This underscores the value of total RNA sequencing for dual host-viral recovery. Furthermore, in the strain associated with specimen M05058, RVA/Nyctophilus geoffroyi- wt/AUS/M05058/1964/G3PX, viral recovery was likely hindered by the use of predominantly *Gammaretrovirus*-targeted probes that were not optimised for RVA. Despite recovering only five segments from specimen M05058, some below confidence thresholds, these partial and cds segments extend the known temporal and taxonomic breadth of bat-associated RVA, representing the earliest molecular evidence of RVA in the genus *Nyctophilus,* which remains underrepresented in global surveillance.

The similarity of strain RVA/Rhinolophus megaphyllus-wt/M24384/1999/G3P[3] to seven other strains across 10 segments, forming the distinct “Au/zoo” constellation (Table 3), demonstrates this genotype constellation has appeared multiple times, in more than one geographical location. The Au/zoo constellation includes multiple bat strains and likely represents a broader bat-associated lineage (Li, et al. 2016; He, et al. 2017; Simsek, et al. 2021). The presence of Au/zoo in Australian *R. megaphyllus* in the 1960s supports a long-standing Australian bat-associated strain. Moreover, the Au/zoo constellation includes human, rodent and equine strains, highlighting its ecological plasticity. Our findings reinforce earlier observations (Simsek, et al. 2021) that bat-associated RVA strains frequently share genotypes with urban wildlife and human lineages, generating strains with zoonotic potential. Given the high frequency of reassortment along with broad host range of RVA strains, such interspecies spillover events are likely to recur (Matthijnssens, Ciarlet, Rahman, et al. 2008).

Understanding the zoonotic potential of rotaviruses of animal origin is critical for surveillance efforts. In Australia, RVA surveillance has primarily focused on livestock as reservoirs (e.g., dairy and beef herds, and piggeries), where zoonotic transmission to humans, particularly children, has been documented (Swiatek, et al. 2010; Cowley, et al. 2013; Genz, et al. 2023). However, the role of bats in this transmission network remains poorly understood. Our data confirms that RVA circulates in Australian bats across the Northern Territory and Queensland, supporting calls for a One Health approach to RVA monitoring that integrates wildlife into public health frameworks.

Globally, bat RVA strains display high genetic diversity, with some strains exhibiting host specificity (e.g., within Pteropodidae) (Simsek, et al. 2021). Continued monitoring in Australian bats, including targeted sampling of *Rhinolophus, Nyctophilus,* and other families, such as Pteropodidae, is needed to map this diversity. Given the established links between ecological stressors (e.g., food shortages) and viral shedding in flying foxes (Becker, et al. 2023), future RVA surveillance should incorporate environmental variables to improve zoonotic risk modelling.

### Spirit vaults as viral archives: unlocking historical viromes from natural history collections

Our recovery of viral and host RNA from a 60-year-old, formalin-fixed microbat specimen advances the growing field of museum metagenomics (Speer, et al. 2022).

This study marks the first characterisation of RVA in an Australian bat and underscores the untapped potential of global vouchered wildlife specimens for studying historical viral diversity. The simultaneous recovery of host and microbial RNA demonstrates the viability of formalin-fixed collections for broad molecular analysis, including eukaryotic, prokaryotic, and viral targets.

Crucially, historical viral RNA has value beyond pathogen detection. It enables viral molecular clock calibration and refinement of evolutionary rate estimates (Worobey, et al. 2008; Duggan, et al. 2016; Duchêne, et al. 2020; Düx, et al. 2020), reveals insights into historical viral taxonomy (Duggan, et al. 2020), and improves our understanding of viral evolution (Taubenberger, et al. 2012). Importantly, our sampling approach was agnostic to symptomatic disease status, in that none of the specimens were selected based on signs of illness, nor did our species selection target members of Chiroptera with known bat-borne viral associations (Wallau, et al. 2023). This approach allows for unbiased recovery of viruses in both symptomatic and asymptomatic hosts. Moreover, formalin fixation inactivates pathogens (Chinabut, et al. 2023; Seeburg, et al. 2023), eliminating biosafety risks and enabling higher-throughput study of high-risk pathogens such as avian influenza, Hendra virus, and lyssaviruses without the need for specialised containment facilities.

By characterising the partial genomes of two historical RVA strains, we demonstrate how biorepositories can resolve phylogenetic gaps, clarify interspecies transmission dynamics, and apply retrospective surveillance, allowing researchers to map virus diversity across time and space. This capacity is particularly valuable for contextualising modern outbreaks and complements real-time modern surveillance in wildlife populations.

Natural history collections are now poised as molecular time capsules, containing critical data for phylogenetics, host-virus coevolution, and emerging infectious disease forecasting. By leveraging chemically fixed wildlife specimens, we can reconstruct the evolutionary dynamics of viruses that shaped, and continue to shape, wildlife and human health. In addition to pandemic preparedness, historical viromes may inform conservation policy, endangered species recovery (where disease plays a role in species decline), landscape-scale microbe mapping, and zoonotic spillover risk models (Thompson Cody, et al. 2021).

This work builds on a growing body of evidence that natural history collections are expanding their original utility as taxonomic archives to become active engines of molecular discovery (Harmon, et al. 2019; Card, et al. 2021; Raxworthy and Smith 2021; Davis and Knapp 2025). Novel scientific contributions of biorepositories, such as unlocking the RNA archives of the spirit vault, are essential for ensuring their continued power to deliver impact for present and future generations of scientists.

When coupled with modern genomic tools, collections transform from molecular graveyards to genomic gold mines which can inform conservation, taxonomy, One Health, and beyond.

## Methods

### Specimen selection

We selected seven specimens from the family Chiroptera, collected between 1964 and 1999, from both the frozen tissue (ethanol-fixed) and whole specimen formalin-fixed collections at the Australian National Wildlife Collection (ANWC) (Table 1).

Specimens ANWC M24834 (*Rhinolophus megaphyllus)* and ANWC M16448 (*Ozimops ridei)*, both collected in the 1990s, were preserved in formalin following dissection and ethanol storage of a subset of internal organs (including liver) at - 80°C. The remaining five specimens, ANWC M05058 (*Nyctophilus geoffroyi)*, ANWC M15981 (*Scotorepens greyii)*, ANWC M16474 (*Hipposideros diadema)*, ANWC M02438 (*Nyctophilus geoffroyi)* and ANWC M19055 (*Mops jobensis)*, were collected between 1964 and 1992 and preserved whole in 10% formalin at collection, before being moved to 70% ethanol.

### RNA extraction

We extracted RNA from approximately 5mm^2^ of liver tissue per specimen. We subsampled tissue from frozen ethanol-preserved samples, and dissected tissue from formalin-preserved whole specimens. We homogenised tissue in 200µL DNA/RNA Shield (Zymo) using sterilised silicon carbide beads (1mm, filled to halfway in an Eppendorf tube) and physical disruption on ice using the Omni Bead Ruptor “skin” protocol (Speed = 7m/s, 30 sec active, 20 sec dwell). We added 400 µg Proteinase K and incubated lysates at 55°C for 4 hours. To pellet the beads and debris, we centrifuged the lysates gently, then processed the supernatant with the Zymo RNA Miniprep kit (Cat# R1057), including DNAse treatment, per manufacturer’s instructions. We assessed RNA fragment length, integrity, and concentration using an Agilent 2100 Bioanalyzer with the Eukaryote Total RNA Pico kit, per manufacturer’s instructions.

### Library preparation and sequencing

We used two different methods of library preparation for the two preservation methods. For ethanol-preserved tissues (ANWC M24834 and M16448), we prepared libraries using the Illumina TruSeq Stranded mRNA kit. For the formalin-preserved tissues (ANWC M05058, M16474, M16474, M02438 and M19055), we prepared total nucleic acid libraries with target enrichment using Twist Bioscience custom RNA probes (details in Supplementary Methods, Supplementary Table 5). The targeted enrichment libraries were sequenced on a Novaseq X plus 10b lane 300 cycle (150 PE). The Illumina TruSeq Stranded mRNA libraries were sequenced on a NovaSeq S4-300 cycle (150 PE). We sequenced all seven libraries at the Australian Genome Research Facility.

### Sequence analysis

#### Quality control

We trimmed adaptor sequences and low-quality bases using Trimmomatic v0.39 (Bolger, et al. 2014) with default settings. We assessed read quality pre- and post-trimming with FastQC v.0.12.1 (Andrews 2010).

#### Identifying viral fragments within libraries

We assembled contiguous sequences *de novo* using both Megahit v1.2.9 (97) and Trinity v2.15.1 (Grabherr, et al. 2011; Li, et al. 2015) with default settings. We similarity-searched the contigs against the NCMI nr database using an unbiased DIAMOND v2.0.15 blastx search via the blast+ package v 2.14.0 (Buchfink, et al. 2021). We considered contigs with viral hits (length > 250bp, percent identity >50%, query cover >50%, *e*-value < 1 x 10^-5^) as viral-like sequences. We mapped raw reads to candidate contigs using the kalign function of the ngskit4b suite version 200218 (https://github.com/kit4b) with options -c25 -l25 -d100-U4 to measure relative abundance, and to fill in any gaps. We calculated mapped read abundance and depth as the total number of reads mapping to the contig divided by the number of total trimmed reads.

#### Viral genome annotation

We annotated RVA contigs using BLAST nucleotide and protein searches. We assigned genotypes to the viral genome segments (n=16) based on nucleotide identity thresholds described in (Matthijnssens, Ciarlet, et al. 2011) and manually annotated open reading frames in Geneious v2024.

### Phylogenetic relationships

We aligned nucleotide sequences (open reading frame regions) for 13 rotavirus segments using MAFFT v7.526 with the L-INS-I algorithm (Katoh, et al. 2002; Katoh and Standley 2013). We inferred maximum likelihood phylogenetic trees from nucleotide alignments, using model testing with 1000 non-parametric bootstrap replicates in IQtree2 v2.2.0.5 (Minh, et al. 2020).

#### Host genome analysis

For specimen ANWC M24834, we mapped reads to the closely related *Rhinolophus affinus* reference genome (CM093080.1) (Zhao, et al. 2024) using STAR v2.7.4a (Dobin, et al. 2013) and visualised the results in Geneious v2024. For other specimens lacking a closely related host genome, we used conserved host genes to estimate host read content (Supplementary Table 3, 4). We calculated host read abundance as the proportion of reads mapped to host chromosomes/genes relative to the total trimmed read count. We also used CCmetagen v1.4.0 (Clausen, et al. 2018; Marcelino, et al. 2020) with the NCBI nt database to estimate overall host RNA representation.

## Acknowledgements

We thank the National Research Collections Australia Digital Operations team, in particular Merinda Campbell and Nicole Fisher, for their photography of ANWC specimens M24834 and M05058. Figures were created with assistance with https://BioRender.com.

## Contributions

Project design: CH, MA Funding: CH, MA, IS, EH. Ethics: MA, CH.

Collection of specimen tissue: MA, CH, TH, CW, IS.

Experimental work: RNA extractions, Twist panel design: MA, Method design: MA, EH, Preservation media survey: EH.

Data analysis: Quality control, Genome assembly, Genome annotation, Phylogenetic trees: AP.

Writing: Original outline; AP, Figures; AP, Editing: AP, EH, CH, CD, TH, CW, IS, MA.

## List of Funding Sources

We acknowledge that this work was funded by CSIRO Environomics Future Science Platform (Grant number R-10011 to CH), CSIRO Heath and Biosecurity Acorn Grant (Grant number R-16935 to MA), CSIRO Early Career Researcher Fellow Program (Grant number R-20592 to CH and AP), National Research Collections Australia Strategic Operating (Grant number R-90579 to CH), and a Department of Agriculture, Fisheries and Forestry Bright Ideas Grant (Grant number R-19122 to MA).

