## Supplementary Materials for "Releasing RNA from formalin: novel 20^th^ century rotavirus alphagastroenteritidis strains identified in Australian microbat voucher specimens"

Supplementary Results

Supplementary Figures 1-10

Supplementary Tables 1-7

Supplementary Methods

### Supplementary Results

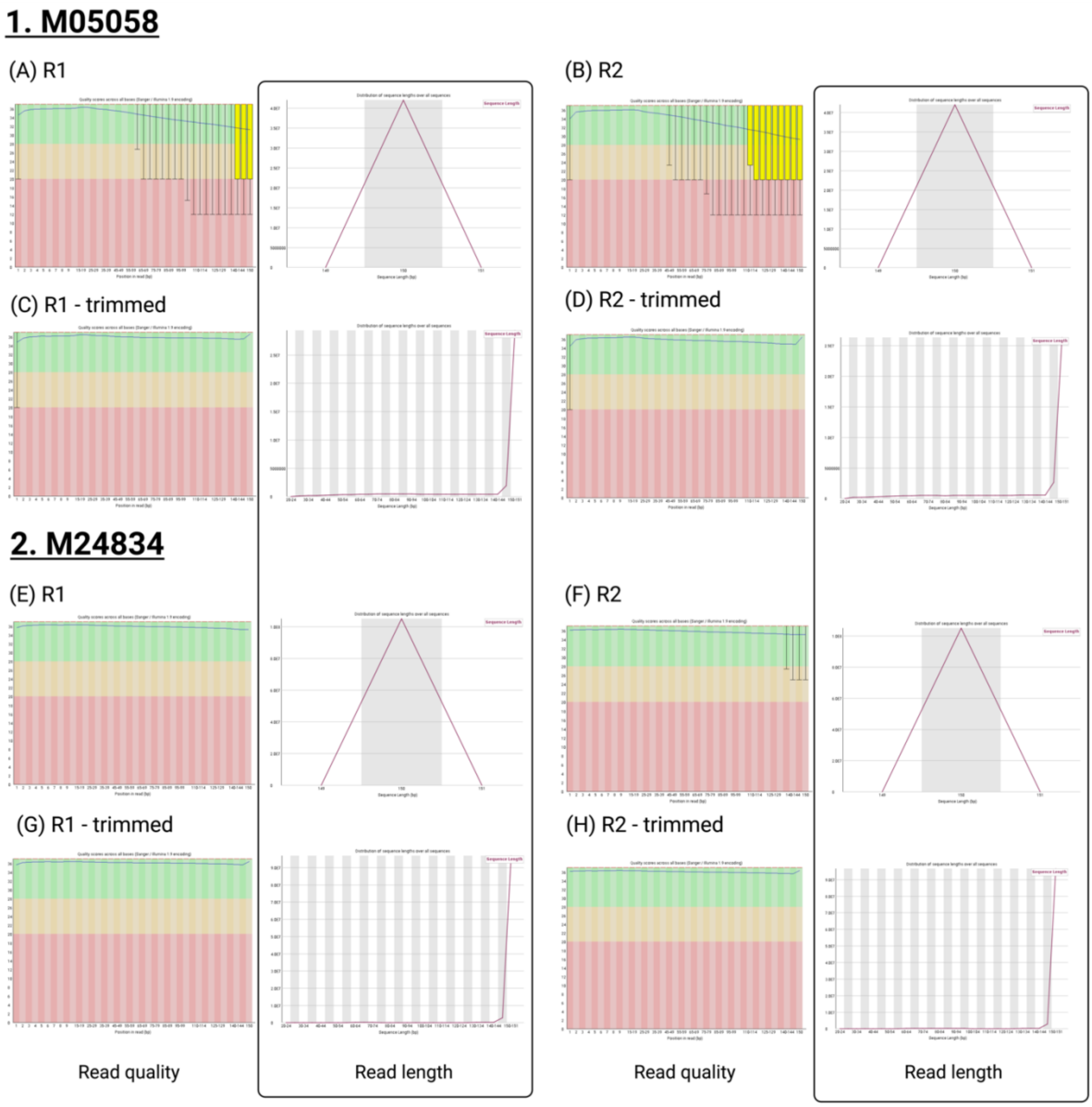


**Supplementary Figure 1.** Read quality analysis determined by FASTQC, showing the read quality plots in the first and third columns, and the read length plots in the second and forth columns, for (1) M05058 and (2) M24834. (A) M05058 read 1 untrimmed. (B) M05058 read 2 untrimmed. (C) M05058 read 1 trimmed. (D) M24834 read 2 trimmed. (E) M24834 read 1 untrimmed. (F) M05058 read 2 untrimmed. (G) M24834 read 1 trimmed. (H) M24834 read 2 trimmed.

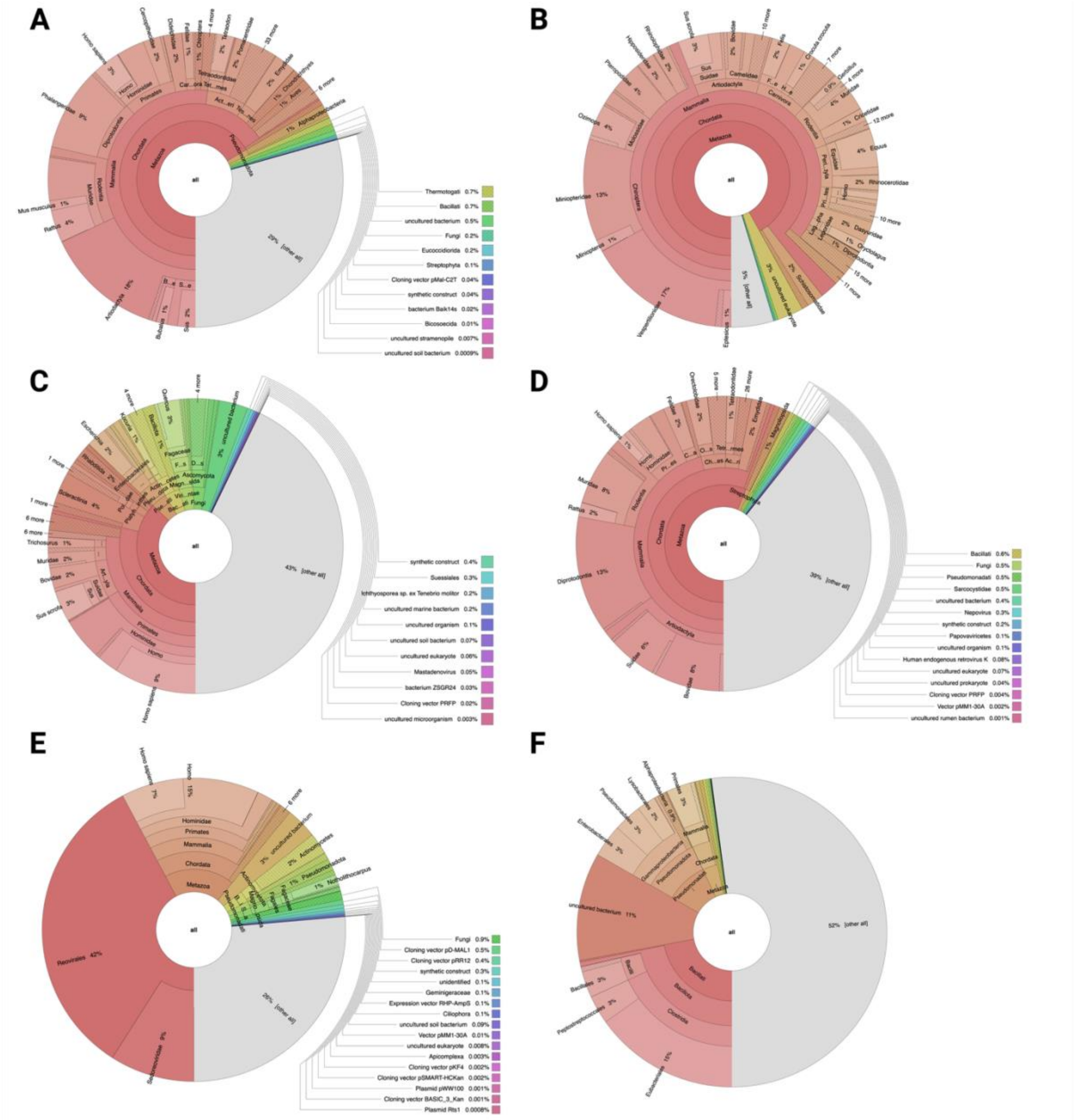


**Supplementary Figure 2**. CCmetagen results for the six remaining specimens examined; M16448, M16474, M02438, M19055, M05058, and M15981. (A) The total trimmed RNA obtained from specimen M16448 has 4% of reads that map to the specimens genus *Ozimops*, more broadly 17% of reads mapping to the family *Vespertillionidae* and 44% mapping to bat order *Chiroptera*, and 86% of reads that map to *Mammalia*. (B) The RNA obtained from specimen M16474 had 0.8% of total trimmed RNA reads map to *Chiroptera* and 66% map to *Mammalia*. (C) The total trimmed RNA obtained from specimen M02438 were broken down into: mapping

0.6% to *Chiroptera*, and mapping 27% to *Mammalia*. (D) Specimen M19055 had total trimmed reads broken down into 0.5% of reads map to *Chiroptera,* 46% of reads map to *Mammalia*, and 39% of reads remain uncharacterised. (E) Total RNA obtained from specimen M05058 has 0.2% total trimmed reads mapping to *Chiroptera*, 46% to *Mammalia*, and 42% to viral family *Reovirales,* and 26% of reads uncharacterised. (F) Specimen M15981 had total trimmed reads broken down into 52% of reads uncharacterised, 22% of reads map to *Bacillati*, 11% of reads map to uncultured bacterium, and 4% of reads map to *Mammalia*.

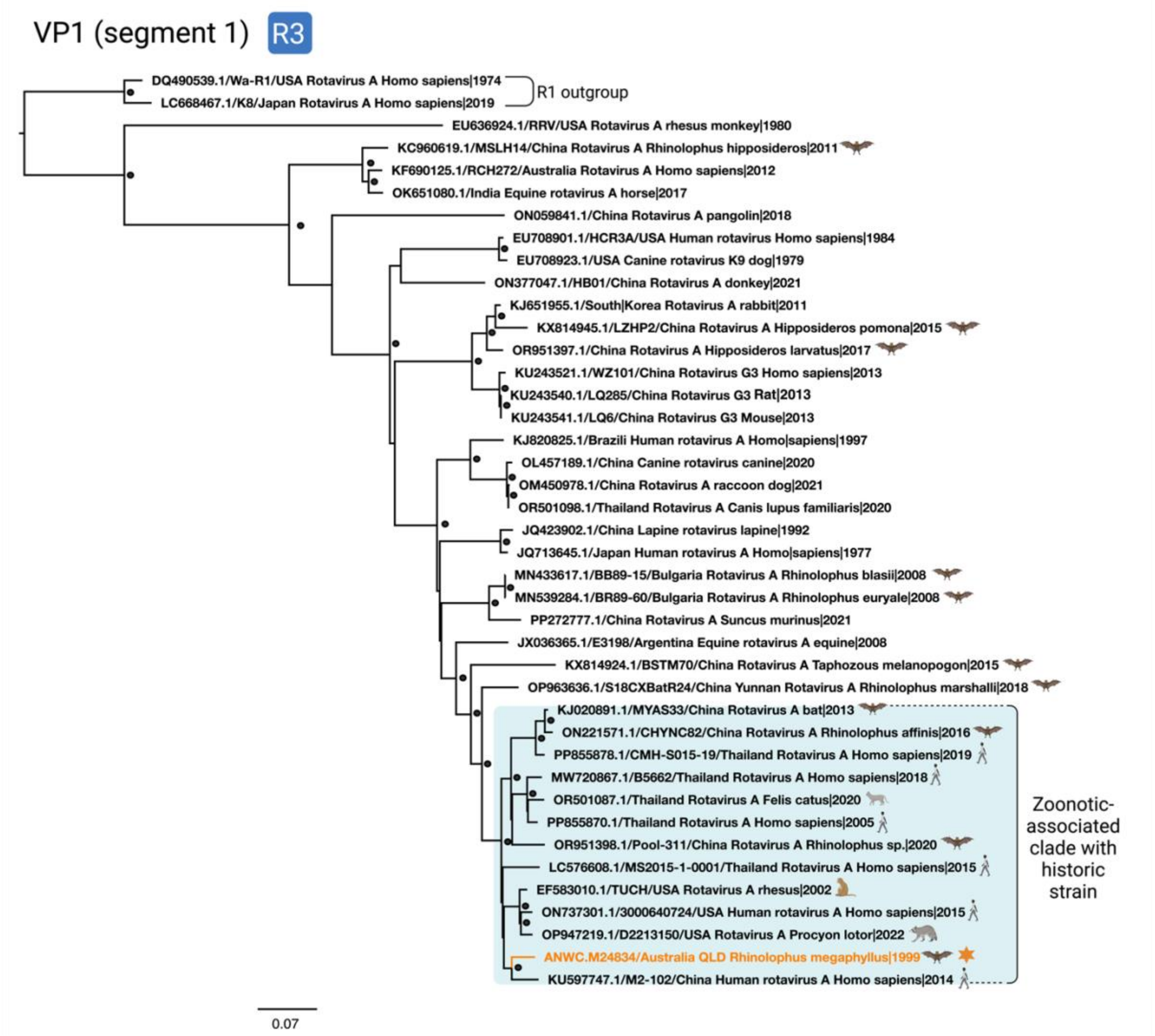


**Supplementary Figure 3.** Phylogenetic tree of RVA segment 1/VP1, containing a cds of VP1 from strain RVA/Rhinolophus megaphyllus-wt/AUS/M24384/1999/G3P[3], along with other partial or whole coding sequences of RVA VP1. genotype R3 (n=40). The novel historical strain has the respective branch highlighted in orange and an orange star next to the tip name. Bat-associated strains have a bat icon next to their tip name. Strains within the same clade as the novel historical strain have their associated animal host icon next to their tip name, and to visualise any inter-species transmission events the historical clade and zoonotic strains are highlighted by a blue box.  Bootstrap replicates > 80% are visualised as filled-in circles at nodes. The scale bar represents nucleotide substitutions per site. For clarity, the phylogeny is rooted at the R1 outgroup.

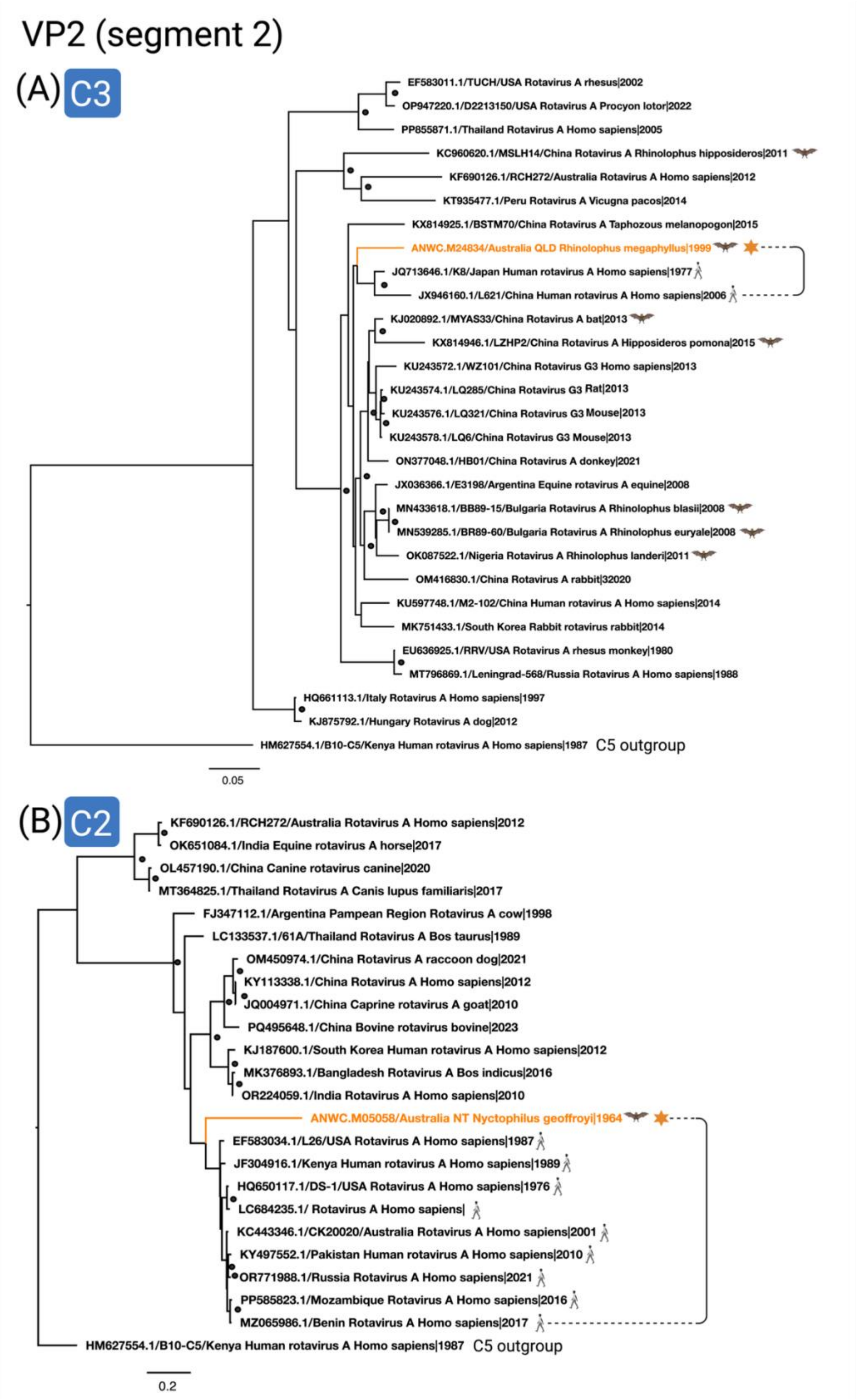
VP2 (segment 2)
(A) C3
EF583011.1/TUCH/USA Rotavirus A rhesus|2002
OP947220.1/D2213150/USA Rotavirus A Procyon lotor|2022
PP855871.1/Thailand Rotavirus A Homo sapiens|2005
KC960620.1/MSLH14/China Rotavirus A Rhinolophus hipposideros|2011
KF690126.1/RCH272/Australia Rotavirus A Homo sapiens|2012
KT935477.1/Peru Rotavirus A Vicugna pacos|2014
KX814925.1/BSTM70/China Rotavirus A Taphozous melanopogon|2015
ANWC.M24834/Australia QLD Rhinolophus megaphyllus|1999
JQ713646.1/K8/Japan Human rotavirus A Homo sapiens|1977
JX946160.1/L621/China Human rotavirus A Homo sapiens|2006
KJ020892.1/MYAS33/China Rotavirus A bat|2013
KX814946.1/LZHP2/China Rotavirus A Hipposideros pomona|2015
KU243572.1/WZ101/China Rotavirus G3 Homo sapiens|2013
KU243574.1/LQ285/China Rotavirus G3 Rat|2013
KU243576.1/LQ321/China Rotavirus G3 Mouse|2013
KU243578.1/LQ6/China Rotavirus G3 Mouse|2013
ON377048.1/HB01/China Rotavirus A donkey|2021
JX036366.1/E3198/Argentina Equine rotavirus A equine|2008
MN433618.1/BB89-15/Bulgaria Rotavirus A Rhinolophus blasii|2008
MN539285.1/BR89-60/Bulgaria Rotavirus A Rhinolophus euryale|2008
OK087522.1/Nigeria Rotavirus A Rhinolophus landeri|2011
OM416830.1/China Rotavirus A rabbit|32020
KU597748.1/M2-102/China Human rotavirus A Homo sapiens|2014
MK751433.1/South Korea Rabbit rotavirus rabbit|2014
EU636925.1/RRV/USA Rotavirus A rhesus monkey|1980
MT796869.1/Leningrad-568/Russia Rotavirus A Homo sapiens|1988
HQ661113.1/Italy Rotavirus A Homo sapiens|1997
KJ875792.1/Hungary Rotavirus A dog|2012
HM627554.1/B10-C5/Kenya Human rotavirus A Homo sapiens|1987
C5 outgroup
0.05
(B) C2
KF690126.1/RCH272/Australia Rotavirus A Homo sapiens|2012
OK651084.1/India Equine rotavirus A horse|2017
OL457190.1/China Canine rotavirus canine|2020
MT364825.1/Thailand Rotavirus A Canis lupus familiaris|2017
FJ347112.1/Argentina Pampean Region Rotavirus A cow|1998
LC133537.1/61A/Thailand Rotavirus A Bos taurus|1989
OM450974.1/China Rotavirus A raccoon dog|2021
KY113338.1/China Rotavirus A Homo sapiens|2012
JQ004971.1/China Caprine rotavirus A goat|2010
PQ495648.1/China Bovine rotavirus bovine|2023
KJ187600.1/South Korea Human rotavirus A Homo sapiens|2012
MK376893.1/Bangladesh Rotavirus A Bos indicus|2016
OR224059.1/India Rotavirus A Homo sapiens|2010
ANWC.M05058/Australia NT Nyctophilus geoffroyi|1964
EF583034.1/L26/USA Rotavirus A Homo sapiens|1987
JF304916.1/Kenya Human rotavirus A Homo sapiens|1989
HQ650117.1/DS-1/USA Rotavirus A Homo sapiens|1976
LC684235.1/ Rotavirus A Homo sapiens|
KC443346.1/CK20020/Australia Rotavirus A Homo sapiens|2001
KY497552.1/Pakistan Human rotavirus A Homo sapiens|2010
OR771988.1/Russia Rotavirus A Homo sapiens|2021
PP585823.1/Mozambique Rotavirus A Homo sapiens|2016
MZ065986.1/Benin Rotavirus A Homo sapiens|2017
HM627554.1/B10-C5/Kenya Human rotavirus A Homo sapiens|1987
C5 outgroup
0.2

**Supplementary Figure 4.** Phylogenetic trees of VP2/segment. The novel historical strain has the respective branch highlighted in orange and an orange star next to the tip name. Bat-associated strains have a bat icon next to their tip name. Strains within the same clade as the novel historical strain have their associated animal host icon next to their tip name, Bootstrap replicates > 80% are visualised as filled-in circles at nodes. The scale bar represents nucleotide substitutions per site. (A) Phylogeny containing the whole coding sequence from strain RVA/Rhinolophus megaphyllus-wt/AUS/M24384/1999/G3P[3], along with other partial or whole coding sequences of RVA VP2, genotype C3 (n=29). For clarity, the phylogeny is rooted at the C5 outgroup. (B) Phylogeny containing the partial sequence from strain RVA/Nyctophilis geoffroyi /AUS/M05058/1964/G3PX, along with other partial or whole coding sequences of RVA VP2, genotype C2 (n=24). For clarity, the phylogeny is rooted at the outgroup C5 strain.

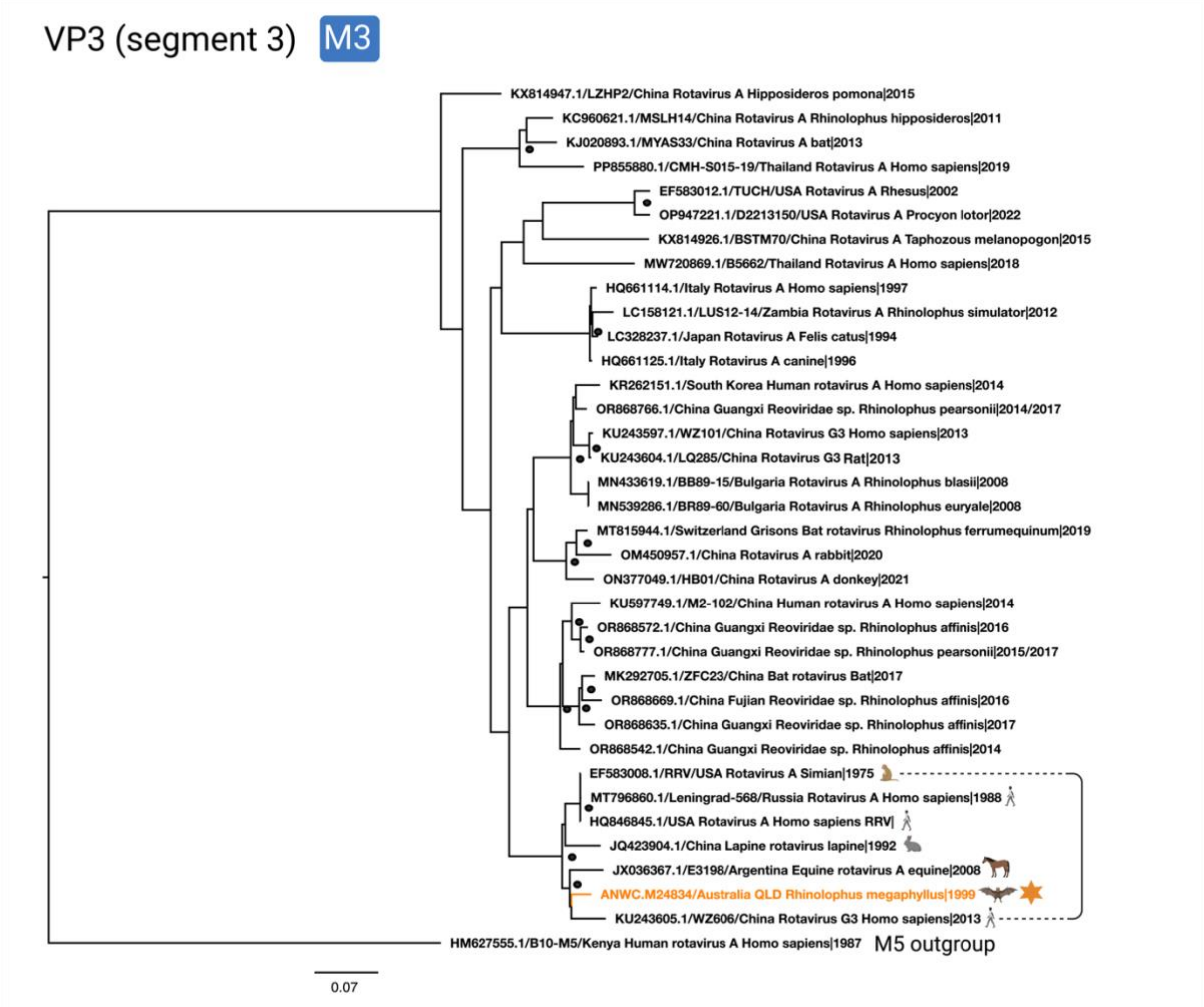


**Supplementary Figure 5.** Phylogenetic tree of VP3/segment 3, containing the whole coding sequence from strain RVA/Rhinolophus megaphyllus-wt/AUS/M24384/1999/G3P[3], along with other partial or whole coding sequences of RVA VP3, genotype M3 (n=37). The novel historical strain has the respective branch highlighted in orange and an orange star next to the tip name. Bat-associated strains have a bat icon next to their tip name. Strains within the same clade as the novel historical strain have their associated animal host icon next to their tip name. Bootstrap replicates > 80% are visualised as filled-in circles at nodes. The scale bar represents nucleotide substitutions per site. For clarity, the phylogeny is rooted at the M5 outgroup.

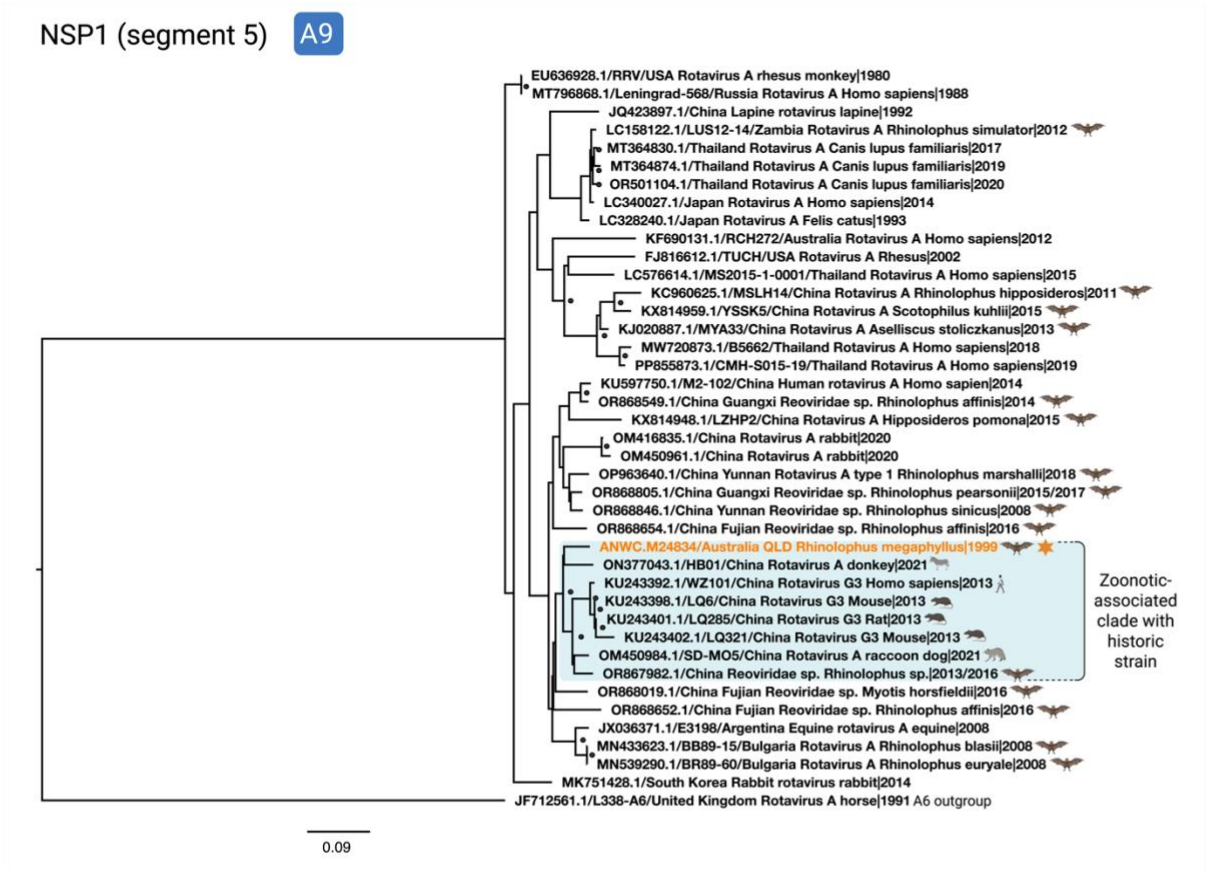


**Supplementary Figure 6.** Phylogenetic tree of NSP1/segment 5, containing the partial sequence from strain RVA/Rhinolophus megaphyllus-wt/AUS/M24384/1999/G3P[3], along with other partial or whole coding sequences of RVA NSP1, genotype A9 (n=41). The novel historical strain has the respective branch highlighted in orange and an orange star next to the tip name. Bat-associated strains have a bat icon next to their tip name. Strains within the same clade as the novel historical strain have their associated animal host icon next to their tip name, to visualise any inter-species transmission events and are highlighted by a blue box. Bootstrap replicates > 80% are visualised as filled-in circles at nodes. The scale bar represents nucleotide substitutions per site. For clarity, the phylogeny is rooted at the A6 outgroup.

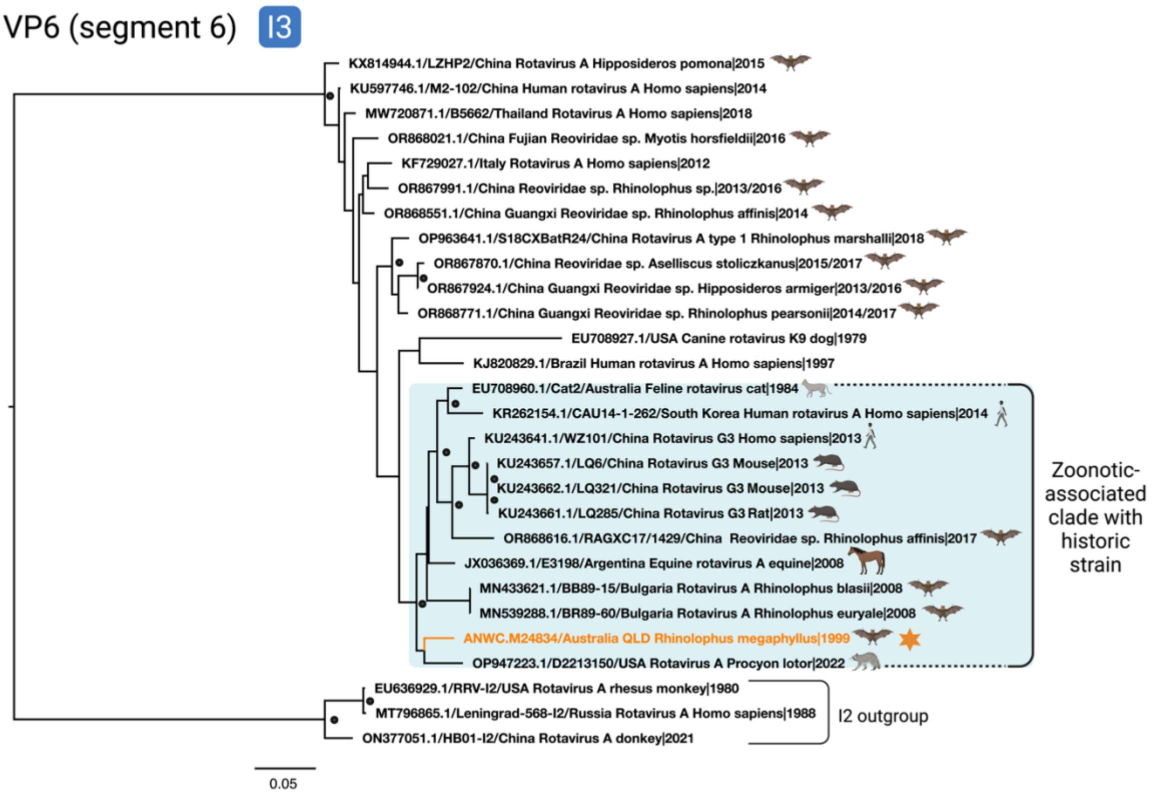


**Supplementary Figure 7.** Phylogenetic tree of VP6/segment 6, containing the partial sequence from strain RVA/Rhinolophus megaphyllus-wt/AUS/M24384/1999/G3P[3], along with other partial or whole coding sequences of RVA VP6, genotype I3 (n=31). The novel historical strain has the respective branch highlighted in orange and an orange star next to the tip name. Bat-associated strains have a bat icon next to their tip name. Strains within the same clade as the novel historical strain have their associated animal host icon next to their tip name, to visualise any inter-species transmission events and are highlighted by a blue box. Bootstrap replicates > 80% are visualised as filled-in circles at nodes. The scale bar represents nucleotide substitutions per site. For clarity, the phylogeny is rooted at the I2 outgroup.

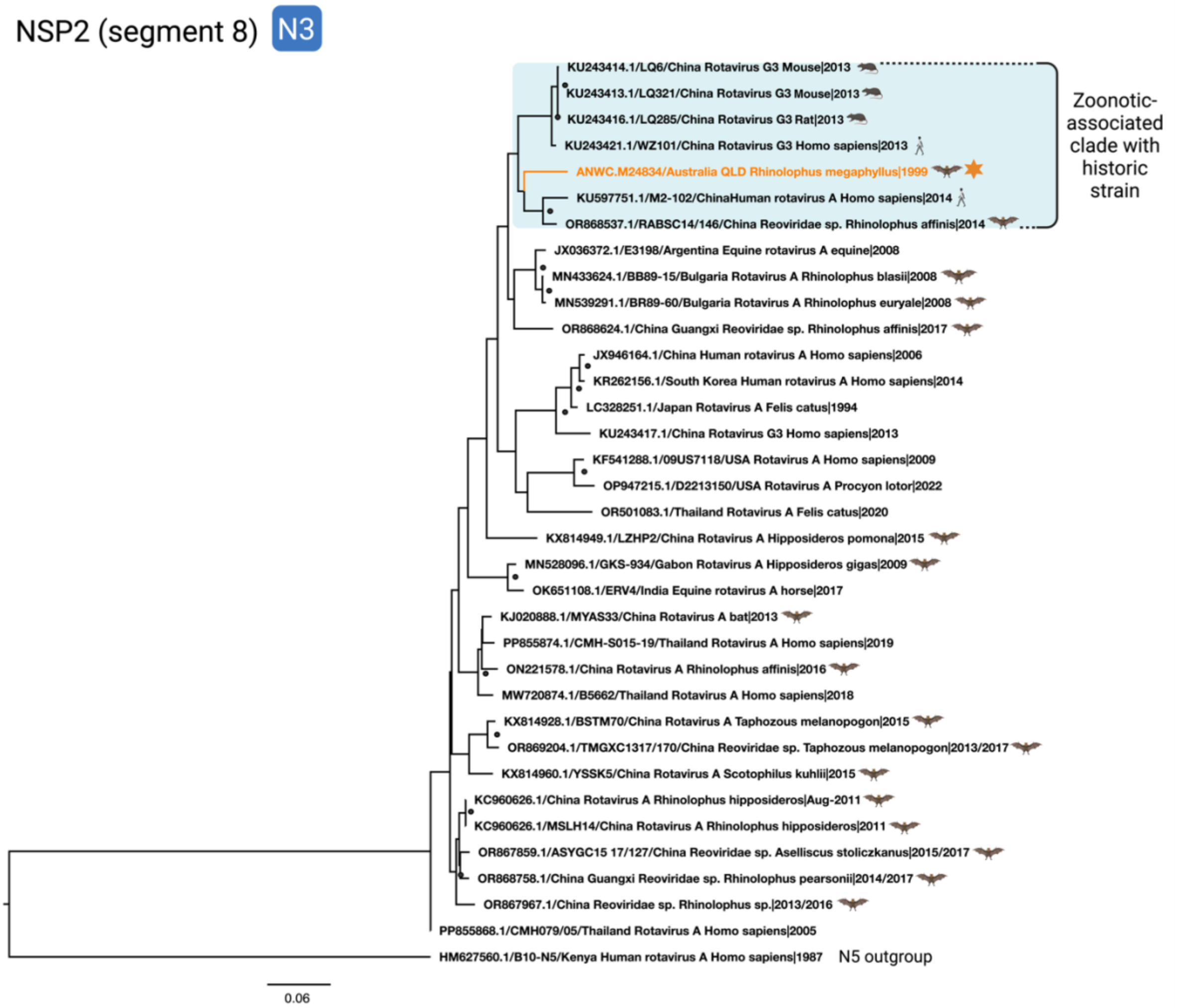


**Supplementary Figure 8.** Phylogenetic tree of NSP2/segment 8, containing the partial sequence from strain RVA/Rhinolophus megaphyllus-wt/AUS/M24384/1999/G3P[3], along with other partial or whole coding sequences of RVA NSP2, genotype N3 (n=35). The novel historical strain has the respective branch highlighted in orange and an orange star next to the tip name. Bat-associated strains have a bat icon next to their tip name. Strains within the same clade as the novel historical strain have their associated animal host icon next to their tip name, to visualise any inter-species transmission events and are highlighted by a blue box. Bootstrap replicates > 80% are visualised as filled-in circles at nodes. The scale bar represents nucleotide substitutions per site. For clarity, the phylogeny is rooted at the N5 outgroup.

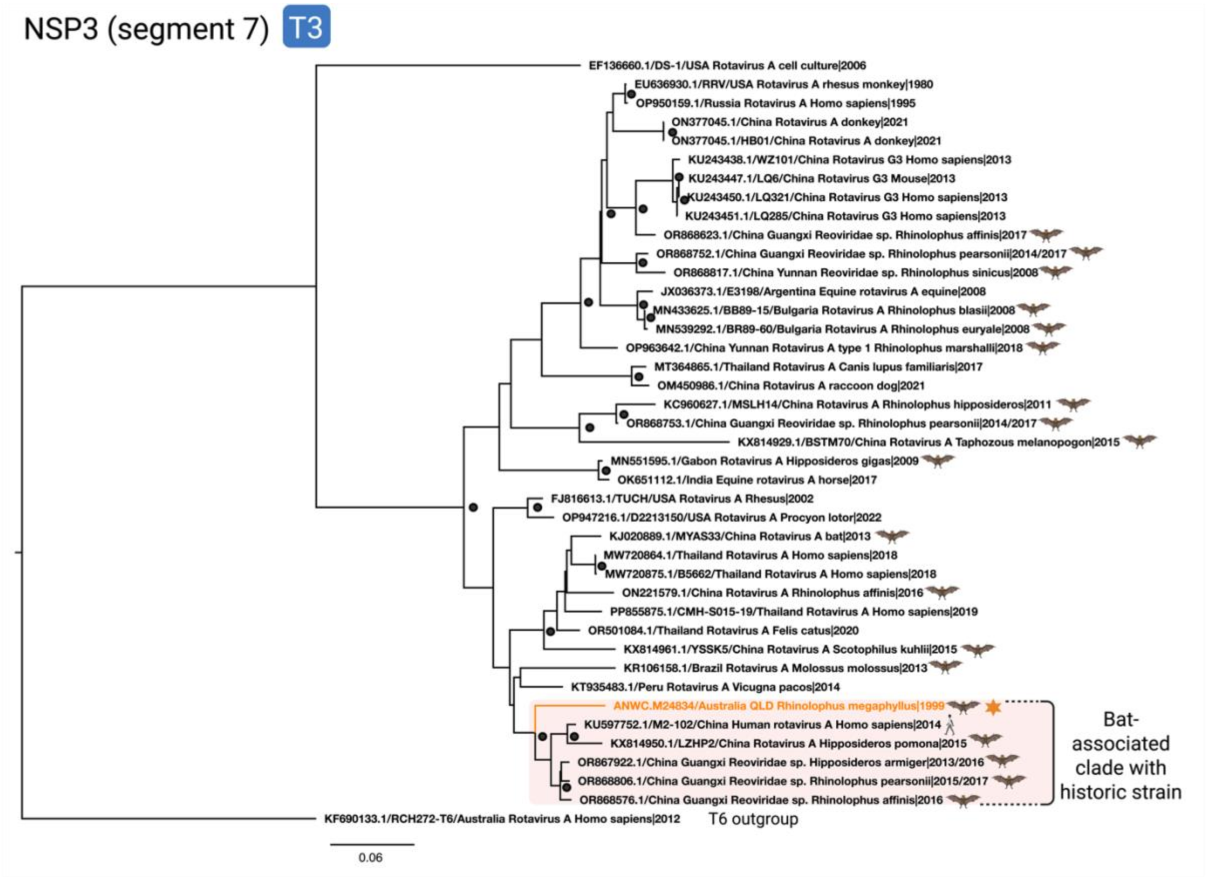


**Supplementary Figure 9.** Phylogenetic tree of NSP3/segment 7, containing the partial sequence from strain RVA/Rhinolophus megaphyllus-wt/AUS/M24384/1999/G3P[3], along with other partial or whole coding sequences of RVA NSP3, genotype T3 (n=41). The novel historical strain has the respective branch highlighted in orange and an orange star next to the tip name. Bat-associated strains have a bat icon next to their tip name. Strains within the same clade as the novel historical strain have their associated animal host icon next to their tip name. To visualise bat-associated clade (majority bat strains) with the historical strain, clade has been highlighted by a red box. Bootstrap replicates > 80% are visualised as filled-in circles at nodes. The scale bar represents nucleotide substitutions per site. For clarity, the phylogeny is rooted at the T6 outgroup.

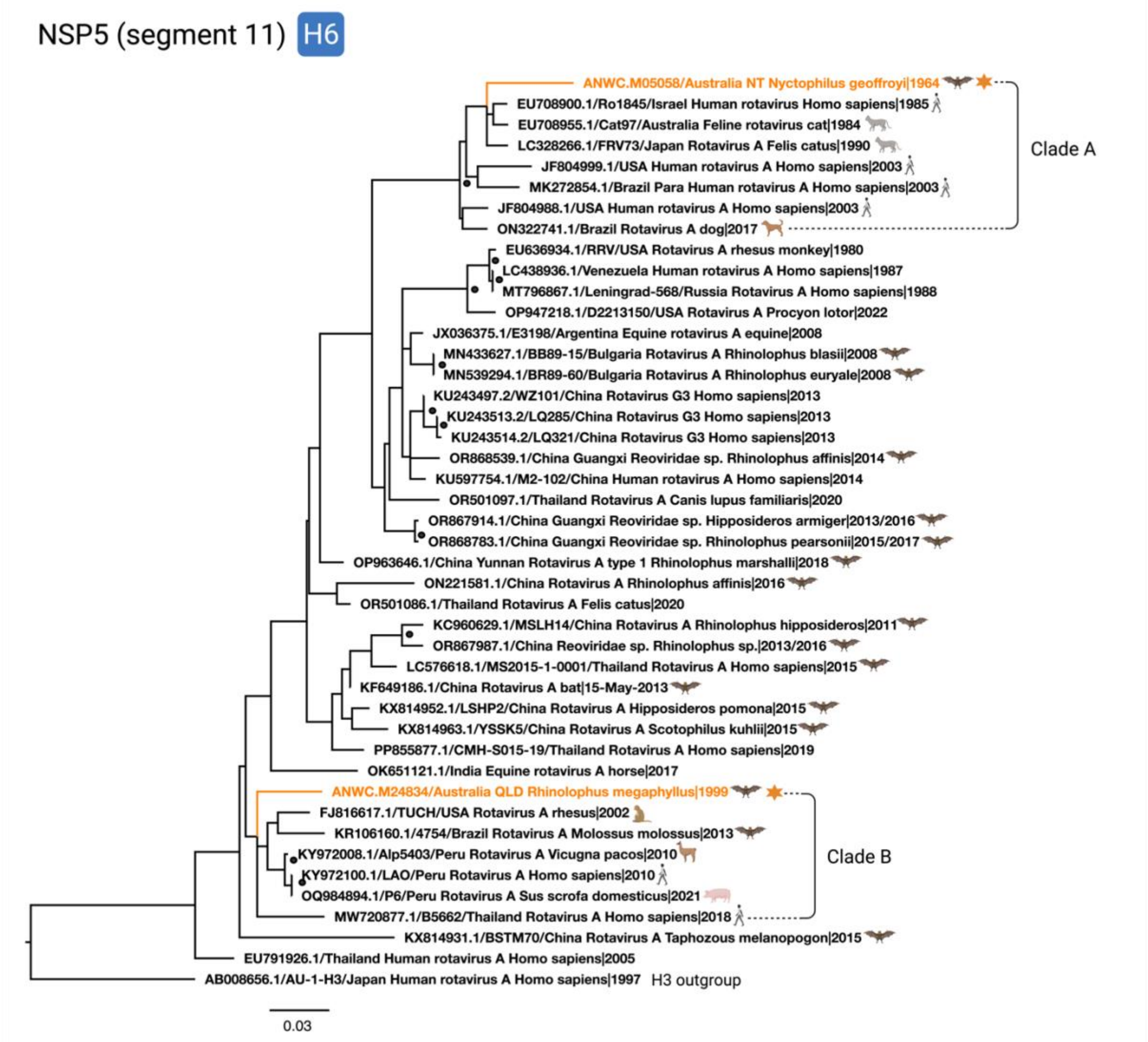


**Supplementary Figure 10.** Phylogenetic tree of NSP5/segment 11. Phylogeny containing the partial sequence from strain RVA/Rhinolophus megaphyllus-wt/AUS/M24384/1999/G3P[3] and the whole sequence from strain RVA/Nyctophilis geoffroyi-wt/AUS/M24384/1964/G3PX, along with other partial or whole coding sequences of RVA NSP5, genotype H6 (n=44). The novel historical strain has the respective branch highlighted in orange and an orange star next to the tip name. Bat-associated strains have a bat icon next to their tip name. Strains within the same clade as the novel historical strain have their associated animal host icon next to their tip name. Bootstrap replicates > 80% are visualised as filled-in circles at nodes.

The scale bar represents nucleotide substitutions per site. For clarity, the phylogeny is rooted at the H3 outgroup.

Supplementary Table 1. FASTQC read quality of sequencing libraries produced by Australian National Wildlife Collection specimens M05058, M24834, M16448, M16474, M02438, M19055, and M15981, pre- and post-trimming.

| Library | | Total sequences | Total Bases (Gbp) | Sequences flagged as poor quality (<20) | %GC | Sequence length | Percent of overrepresented sequences* |
|---|---|---|---|---|---|---|---|
| M05058 | R1 | 41979768 | 6.2 | 0 | 58 | 150 | 95.1 |
| | R2 | 41979768 | 6.2 | 0 | 58 | 150 | 95.1 |
| M05058 trimmed | R1 | 40429673 | 5.5 | 0 | 57 | 25-150 | 96.2 |
| | R2 | 40429673 | 5.3 | 0 | 57 | 25-150 | 96.2 |
| M24834 | R1 | 104862203 | 15.7 | 0 | 50 | 150 | 61.3 |
| | R2 | 104862203 | 15.7 | 0 | 50 | 150 | 57.8 |
| M24834 trimmed | R1 | 103918974 | 15.3 | 0 | 50 | 25-150 | 72.2 |
| | R2 | 103918974 | 15.2 | 0 | 50 | 25-150 | 68 |
| M16448 | R1 | 109970546 | 16.4 | 0 | 50 | 150 | 78.3 |
| | R2 | 109970546 | 16.4 | 0 | 50 | 150 | 78.3 |
| M16448 trimmed | R1 | 109029133 | 16 | 0 | 50 | 25-150 | 77.5 |
| | R2 | 109029133 | 16 | 0 | 50 | 25-150 | 77.5 |
| M16474 | R1 | 437782014 | 6.5 | 0 | 57 | 150 | 97 |
| | R2 | 437782014 | 6.5 | 0 | 57 | 150 | 96.1 |
| M16474 trimmed | R1 | 42550727 | 6.1 | 0 | 57 | 25-150 | 97.5 |
| | R2 | 42550727 | 6.1 | 0 | 57 | 25-150 | 97.5 |
| M02438 | R1 | 28407629 | 4.2 | 0 | 58 | 150 | 96.4 |
| | R2 | 28407629 | 4.2 | 0 | 58 | 150 | 94.5 |
| M02438 trimmed | R1 | 27401174 | 3.8 | 0 | 58 | 25-250 | 97 |
| | R2 | 27401174 | 3.7 | 0 | 58 | 25-150 | 96.6 |
| M19055 | R1 | 35308523 | 5.2 | 0 | 59 | 150 | 96 |
| | R2 | 35308523 | 5.2 | 0 | 59 | 150 | 94.7 |
| M19055 trimmed | R1 | 34155912 | 4.8 | 0 | 58 | 25-150 | 96.5 |
| | R2 | 34155912 | 4.7 | 0 | 58 | 25-150 | 96.3 |
| M15981 | R1 | 60231089 | 9 | 0 | 57 | 150 | 96.2 |
| | R2 | 60231089 | 9 | 0 | 57 | 150 | 95.1 |
| M15981 trimmed | R1 | 58102523 | 8.3 | 0 | 57 | 25-150 | 96.8 |
| | R2 | 58102523 | 8.3 | 0 | 57 | 25-150 | 96.8 |

*less than 30bp, kmer dimers or highly repetitive tandem repeats

Supplementary Table 2. Read abundance of candidate viral contigs from samples M05058 and M24834 that demonstrated similarity to Rotavirus alphagastroenteritidis at the nucleotide or protein level through BLAST searching.

| Sample | Segment and protein | Reads mapped | Read abundance (%)* |
|---|---|---|---|
| M05058 | Segment 9 (VP7), partial | 8,763 | 0.02 |
| | Segment 4 (VP4), partial | 8,098 | 0.02 |
| | Segment 6 (VP6), partial | 78,935 | 0.2 |
| | Segment 2 (VP2), partial | 22,084 | 0.05 |
| | Segment 11 (NSP5), partial | 2,561,774 | 6.3 |
| M24384 | Segment 9 (VP7), cds | 127 | 0.0001 |
| | Segment 4 (VP4), partial | 41 | 0.00004 |
| | Segment 6 (VP6), partial | 24 | 0.00002 |
| | Segment 1 (VP1), partial | 201 | 0.0002 |
| | Segment 11 (NSP5), partial | 10 | 0.00001 |
| | Segment 7 (NSP3), partial | 125 | 0.0001 |
| | Segment 6 (NSP2), partial | 137 | 0.0001 |
| | Segment 5 (NSP1), partial | 130 | 0.0001 |
| | Segment 3 (VP3), cds | 427 | 0.0004 |
| | Segment 2 (VP2), cds | 386 | 0.0004 |

*Measured as reads mapped to final contig, divided by total number of reads in library, calculated as a percentage of 100. Total library reads post trimming equal: M05058 n= 40,429,673, M24834 n=103,918,974.

Supplementary Table 3. Read abundance of M24834 mapping to whole genome sequence of *Rhinolophus affinis* (CM093080.1).

| Sample | Segment | Sequence length | Reads mapped* | Read abundance (%) |
|---|---|---|---|---|
| ANWC M24834 | Chromosome 30 | 16,973,988 | 750,327 | 0.7 |
| | Chromosome 29 | 21,155,509 | 1,687,826 | 1.6 |
| | Chromosome 28 | 26,108,459 | 2,088,524 | 2 |
| | Chromosome 27 | 27,864,874 | 1,398,679 | 1.3 |
| | Chromosome 26 | 29,305,267 | 1,406,162 | 1.4 |
| | Chromosome 25 | 30,169,554 | 682,795 | 0.7 |
| | Chromosome 24 | 32,188,157 | 1,078,497 | 1 |
| | Chromosome 23 | 41,660,319 | 1,307,511 | 1.3 |
| | Chromosome 22 | 42,700,839 | 2,403,515 | 2.3 |
| | Chromosome 21 | 45, 518,199 | 2,417,599 | 2.3 |
| | Chromosome 20 | 46,420,119 | 2,052,699 | 2 |
| | Chromosome 19 | 52,715,897 | 3,573,945 | 3.4 |
| | Chromosome 18 | 57,560,827 | 11,634,868 | 11.2 |
| | Chromosome 17 | 59,592,074 | 1,548,774 | 1.5 |
| | Chromosome 16 | 60,187,402 | 1,489,307 | 1.4 |
| | Chromosome 15 | 64,142,221 | 3,189,978 | 3.1 |
| | Chromosome 14 | 67,976,831 | 2,768,894 | 2.7 |
| | Chromosome 13 | 68,861,389 | 2,771,234 | 2.7 |
| | Chromosome 12 | 79,013,317 | 3,315,721 | 3.2 |
| | Chromosome 11 | 85,492,230 | 3,468,615 | 3.3 |
| | Chromosome 10 | 91.599.250 | 6,545,408 | 6.3 |
| | Chromosome 9 | 93,989,656 | 4,670,667 | 4.5 |
| | Chromosome 8 | 95,957,452 | 4,172,211 | 4 |
| | Chromosome 7 | 101,258,431 | 2,804,232 | 2.7 |
| | Chromosome 6 | 102,801,965 | 3,263,647 | 3.1 |
| | Chromosome 5 | 103,304,885 | 10,853,297 | 10.4 |
| | Chromosome 4 | 105,875,891 | 2,355,951 | 2.2 |
| | Chromosome 3 | 110,689,453 | 3,376,701 | 3.2 |
| | Chromosome 2 | 112,076,870 | 3,884,626 | 3.7 |
| | Chromosome 1 | 112,511,327 | 5,290,668 | 5.1 |
| | Chromosome X | 121,107,652 | 2,841,928 | 2.7 |
| | Mitochondrial | 17,034 | 18,909 | 0.02 |
| | Total percentage | | | |
| *Not unique mapping, includes repetitive mapping | | | | |

Supplementary Table 4. Read abundance of host-like fragments from samples M16448, M16474, M02438, M05058, M15981, and M19055.

| Sample | Species | Accession number | Gene name | Sequence length (bp) | Reads mapped | Read abundance (%)* |
|---|---|---|---|---|---|---|
| M16448 | *Ozimops ridei* | KJ588342.1 | NADH dehydrogenase subunit 2, partial. | 759 | 31,026 | 0.03 |
| M16474 | *Hipposideros diadema* | KU834053.1 | Hdl_33 olfactory receptor family 52 (OR52) gene, partial cds. | 723 | N/A | N/A |
| M02438 | *Nyctophilus geoffroyi* | MT246225.1 | Cytochrome C oxidase subunit I (COX1) gene, partial, mitochondrial | 658 | 3,706 | 0.01 |
| M05058 | *Nyctophilus geoffroyi* | MT246225.1 | Cytochrome C oxidase subunit I (COX1) gene, partial, mitochondrial | 658 | 4,432 | 0.01 |
| M15981 | *Scotomanes ornatus* | HM561701.1 | Dentin matrix acidic phosphoprotein I (DMP1) gene, partial cds | 951 | N/A | N/A |
| M19055 | *Mops jobensis* | AY591331.1 | Cytochrome B gene, partial, mitochondrial | 423 | N/A | N/A |
| *Measured as reads mapped to gene, divided by total number of trimmed reads in library, calculated as a percentage of 100. Total library reads post trimming equal: M05058 n= 40,429,673, M24834 n=103,918,97, M1648= 109,029,133, M02438= 27,401,174 | | | | | | |

**Supplementary Table 5. Genomes used to design custom Twist panel.**

| Genome reference ID | Genome name |
| --- | --- |
| KF572483.1 | Melomys burtoni retrovirus isolate BRME001 polymerase (Pol) gene, partial cds |
| MN413610.1 | Hervey pteropid gammaretrovirus isolate HPG, complete genome |
| KF572486.1 | Melomys burtoni retrovirus isolate BRME004 envelope glycoprotein (Env) gene, partial cds |
| KF572485.1 | Melomys burtoni retrovirus isolate BRME003 envelope glycoprotein (Env) gene, partial cds |
| KF572484.1 | Melomys burtoni retrovirus isolate BRME002 envelope glycoprotein (Env) gene, partial cds |
| MT062420.1 | Achimota pararubulavirus 3 isolate U72, complete genome |
| NC_034401.1 | Quezon virus clone MT1720/1657 segment L, complete sequence |
| N/A | PAdV_B03 adenovirus |
| N/A | PAdV8A_Hb7B adenovirus |

Supplementary Table 6. Megablast results for each contig, listing top result from non-metagenomic studies.

| Sample | Segment | Closest known relative (accession) | Closest known relative (strain/isolate) | Identified in host | Percent Identity (%) | Query Cover (%) | E value |
|---|---|---|---|---|---|---|---|
| M05058 | Segment 2 (VP2), partial | EF583034.1 | RVA/Human-tc/PHL/L26/1987/G12P[4] | Lab-adapted | 83.71 | 98 | 1e-75 |
| | Segment 4 (VP4), partial | LC328211.1 | RVA/Cat-tc/JPN/FRV72/1990/G3P[3] | *Felis catus* | 79.91 | 91 | 5e-33 |
| | Segment 6 (VP6), partial | LC228363.1 | RVA/Human-wt/JPN/K-21-16/2016/G2P[4] | *Homo sapiens* | 80.85 | 100 | 7e-141 |
| | Segment 7 (NSP2), partial | PP861903.1 | RVA/Human-wt/CHN/Fuzhou21-41/2021/G9P[8] | *Homo sapiens* | 100 | 49 | 1e-21 |
| | Segment 9 (VP7), partial* | KT250942.1 | RVA/ALP/Peru/3386-10/2010 | *Lama pacos* | 85.04 | 90 | 5e-64 |
| | Segment 9 (VP7), partial* | KY418050.1 | RVA/Human-wt/TWN/105-606-D039/2016/G3P8 | *Homo sapiens* | 85.66 | 76 | 3e-66 |
| | Segment 11 (NSP5), partial | EU708955.1 | RVA/Cat/AUS/Cat97/G3P[3] | *Felis catus* | 96.92 | 99 | 0.0 |
| M24834 | Segment 1 (VP1), partial | EF583010.1 | RVA/Rhesus-tc/USA/TUCH/2002/G3P[24] | *Macaca mulatta*, lab-adapted | 93.93 | 100 | 0.0 |
| | Segment 2 (VP2), cds | JQ713646.1 | RVA/Human-tc/JPN/K8/1977/G1P[9] | *Homo sapiens* | 93.81 | 100 | 0.0 |
| | Segment 3 (VP3), cds | EU636926.1 | RVA/Simian-tc/USA/RRV/1975/G3P[3] | Macaca mulatta | 95.57 | 100 | 0.0 |
| | Segment 4 (VP4), partial | KU597745.1 | RVA/Human-wt/CHN/M2-102/2014/G3P[3] | *Homo sapiens* | 90.67% | 100 | 4e-166 |
| | Segment 5 (NSP1), partial | ON377043.1 | RVA/Donkey-wt/HB01/China/2021/G3P12 | *Equus asinus* | 92.97 | 100 | 0.0 |
| | Segment 6 (VP6), partial | LC328220.1 | RVA/Cat-tc/JPN/FRV317/1994/G3P[9] | *Felis catus* | 95.04 | 100 | 0.0 |
| | Segment 7 (NSP2), partial | KU243421.1 | RVA/Human/CHN/WZ101/2013 | *Homo sapiens* | 92.86 | 100 | 0.0 |
| | Segment 8 (NSP3), partial | KU597752.1 | RVA/Human-wt/CHN/M2-102/2014/G3P[3] | *Homo sapiens* | 94.53 | 100 | 0.0 |
| | Segment 9 (VP7), cds | MN433622.1 | RVA/Bat-wt/BGR/BB89-15/2008/G3P3 | *Rhinolophus blasii* | 95.92 | 100 | 0.0 |
| | Segment 11 (NSP5), partial | FJ816617.1 | RVA/Rhesus-tc/USA/TUCH/2002/G3P[24] | Unspecified | 95.82 | 100 | 0.0 |

*Separate megablast hits for each contig

**Supplementary Table 7.** Best-fit DNA models used in phylogenetic analysis, chosen according to Bayesian Information Criterion by ModelFinder in IQ-TREE2.

| **Phylogenic tree** | **Model** |
|---|---|
| Segment 9 (VP7) | TIM2+F+G4 |
| Segment 1 (VP1) | TIM3+F+G4 |
| Segment 2 (VP2) – C2 | GTR+F+G4 |
| Segment 2 (VP2) – C3 | TIM3+F+G4 |
| Segment 3 (VP3) | TIM3+F+G4 |
| Segment 4 (VP4) | TPMu+F+G4 |
| Segment 5 (NSP1) | TIM3+F+I+G4 |
| Segment 6 (VP6) | TIM2+F+G4 |
| Segment 7 (NSP2) | TIM3+F+G4 |
| Segment 8 (NSP3) | TIM3+F+G4 |
| Segment 11 (NSP5) | HKY+F+I+G4 |

### Supplementary Results

**Relationships to previously identified RVA strains (continued)**

The complete coding sequence of VP7 (genotype G3) characterised from RVA/Rhinolophus megaphylls/AUS/M24384/1999/G3P[3] clustered within a clade of strains that have previous evidence of zoonotic or inter-species transmission. Specifically, the novel historical strain was found to be closely related to two unique human strains of RVA with evidence of cross-species transmission (Figure 2b, blue box). RVA/Human-wt/CHN/WZ101/2013/G3P[3] is potentially zoonotic and associated with a group of endemic rodent strains (Li, et al. 2016), and RVA/Human-wt/DOM/Dom114/2007/G3PX was isolated from a child hospitalised with gastroenteritis in the Dominican Republic in 2007, but shared high genetic similarity to Bulgarian bat RVA strains RVA/Bat-wt/BGR/BB89-15/2008/G3P[3] and RVA/Bat-wt/BGR/BR89-60/2008/G3P[3] (>= 97% nucleotide identity) (Bourdett-Stanziola, et al. 2021), suggesting interspecies transmission events.

Other closely related animal strains include the rodent strains linked to RVA/Human-wt/CHN/WZ101/2013/G3P[3], suggested to be zoonotic, RVA/Rat-wt/CHN/LQ285/2013/G3P[3] identified in *Rattus losea* (lesser ricefield rat), and RVA/Mouse/CHNLQ6/2013/G3P[3] and RVA/Mouse/CHNLQ321/G3P[3] both identified in *Alexandromys fortis* (reed vole). Two bovine strains from Spain and China were also clustered within the zoonotic-associated clade, including strain RVA/Donkey-wt/CHN/HB01/2021/G3P[12]. The importance of zoonotic risk in bovine strains, along with the impact of rotavirus in diarrheic calves on productivity loss, has previously been demonstrated (Benito, et al. 2020; Seid, et al. 2020). The other two strains included in the clade were detected in bats from Bulgarian caves in 2008 (Simsek, et al. 2021). The strains, RVA/Bat-wt/BGR/BB89-15/2008/G3P[3] sampled from *Rhinolophus blaslii* (Blasius's horseshoe bat) and RVA/Bat-wt/BGR/BR89-60/2008/G3P[3] sampled from *Rhinolophus euryale* (Mediterranean horseshoe bat), demonstrate high sequence and genotype constellation similarity to RVA/Rhinolophus megaphylls/AUS/M24384/1999/G3P[3]) (Table 3, Supplementary Table 6).

The partial VP4 (genotype P[3]) sequence characterised from RVA/Rhinolophus megaphylls/AUS/M24384/1999/G3P[3] clusters within a bat-associated clade in the phylogeny, with singular human strain RVA/Human-wt/CHN/M2-102/2014/G3P[3] (Figure 2c, red box). The RVA/Human-wt/CHN/M2-102/2014/G3P[3] strain is associated with suspected wildlife reassortment events (Dong, et al. 2016), Furthermore, bat strain RVA/Bat-wt/CHN/LZHP2/2015/G3P[3]. detected in *Hipposideros pomona* (Pomona roundleaf bat) (He, et al. 2017), was highly similar in both nucleotide identity and genotype constellation to strain M2-102. Previous studies have suggested an interspecies transmission link, as strain M2-102 was detected in a 3-year-old child with diarrhea several years prior (Dong, et al. 2016), just 120km from where the LZHP2 was detected (56). The closest relative is a *Rhinolophus*-associated strain RVA/Bat-wt/CH/Rhi_fer/2019 collected from *R. ferrumequinum* in Switzerland (Hardmeier, et al. 2021).

Other segments of RVA/Rhinolophus megaphylls/AUS/M24384/1999/G3P[3] are closely related to respective segments of strain M2-102. For example, in the VP1 phylogenetic tree, the concatenated sequence of RVA/Rhinolophus megaphylls/AUS/M24384/1999/G3P[3] VP1 (genotype R3) is closely related to strain RVA/Human-wt/CHN/M2-102/2014/G3P[3] (Supplementary Figure 3). Bat strains related to the historical strain include RVA/Bat-tc/CHN/MYAS33/2013/G3P[10], detected in an unspecified bat from China in 2013, which also shares a highly similar genotype constellation (Table 3), and two strains isolated from Chinese *Rhinolophus sp.* RVA/Bat-tc/CHN/CHYNC82/2016/G3P3 and isolate Pool-311. The phylogenetic relationships between bat strains and the human strain found in the bat-associated VP1 clade, RVA/Human-wt/THA/CMH-S015-19/2019, suggest interspecies transmission of RVA between bats and humans, and contribute to evidence that the evolution of human RVA strains is closely interwoven with zoonotic RVA strains (Jampanil, et al. 2024).

Within the NSP1 phylogeny, the RVA/Rhinolophus megaphylls/AUS/M24384/1999/G3P[3] NSP1 partial sequence (genotype A9) clustered with potentially zoonotic RVA/Human-wt/CHN/WZ101/2013/G3P[3] and associated endemic rodent strains (Li, et al. 2016), and is closely related to bovine strain RVA/CHN/Donkey-wt/HB01/2021/G3P[12] (Supplementary Figure 6). The partial VP6 (genotype I3) sequence from RVA/Rhinolophus megaphylls/AUS/M24384/1999/G3P[3] clusters within the same clade as potentially zoonotic RVA/Human-wt/CHN/WZ101/2013/G3P[3] and associated endemic rodent strains (Li, et al. 2016), the two bat strains from Bulgaria, and an unusual equine strain RVA/Horse-wt/ARG/E3198/2008/G3P[3] (Supplementary Figure 7). The E3198 strain has evidence of multiple interspecies transmission and/or reassortment events (Miño, et al. 2013). The closest relative to RVA/Rhinolophus megaphylls/AUS/M24384/1999/G3P[3] within the VP6 phylogeny was RVA/Racoon/USA/D2213150/2022/G3P[3], isolated from a raccoon (*Procyon lotor*).

In the NSP2 phylogeny, the RVA/Rhinolophus megaphylls/AUS/M24384/1999/G3P[3] NSP2 partial sequence (genotype N3) is closely related to RVA/Human-wt/CHN/M2-102/2014/G3P[3], and potentially zoonotic RVA/Human-wt/CHN/WZ101/2013/G3P[3] and associated endemic rodent strains (Li, et al. 2016) Supplementary Figure 8). In the NSP3 phylogenetic tree, the RVA/Rhinolophus megaphylls/AUS/M24384/1999/G3P[3] NSP3 partial sequence (genotype T3) falls basal to strain RVA/Human-wt/CHN/M2-102/2014, along with four bat strains forming a bat-associated clade (Supplementary Figure 9).

Within the VP3 phylogeny (Supplementary Figure 5), the VP3 (genotype M3) cds from RVA/Rhinolophus megaphylls/AUS/M24384/1999/G3P[3] is closely related to a strain indicated to be the product of cross-species transmission, RVA/Human-wt/WZ606/2013 (Li, et al. 2016), and to potentially zoonotic and/or reassortment-associated equine strain E3198 (Miño, et al. 2013). The VP2 cds (genotype C3) isolated from RVA/Rhinolophus megaphylls/AUS/M24384/1999/G3P[3] was closely related to two human strains, RVA/Human-tc/JPN/K8/1977/G1P[9] from Japan in 1977 and strain RVA/Human-tc/CHN/L621/2006/G3P[9] from China in 2006 (Supplementary Figure 4a). The partial VP2 (genotype C2) sequence from RVA/Nyctophilus geoffroyi-wt/AUS/M05058/1964/G3PX clusters basal to a DS-1-like human clade within the VP2 phylogeny (Supplementary Figure 4b). Of note, no other bat strains are known to have a C2 genotype. Australian strain RVA/Human-wt/RCH272/2012/G3P[14] is distantly related. Within the NSP5 phylogeny

(Supplementary Figure 10), both RVA/Rhinolophus megaphyllus-wt/AUS/M24384/1999/G3P[3] and RVA/Nyctophilus geoffroyi-wt/AUS/M05058/1964/G3PX NSP5 partial sequences (genotype H6) clustered within multi-species clades.

### Supplementary methods

#### RNA extraction, library preparation and sequencing

RNA samples were quantified and diluted with nuclease-free water to a concentration of 3.3 ng/µL in a 15 µL volume. For samples with RNA concentrations below 3.3 ng/µL (4 out of the 22 samples), the full 15 µL of the sample was used for cDNA synthesis.

In consultation with Australian bat virologists, we identified that the Twist Comprehensive Viral Research panel (#103545) lacked coverage for several known bat viruses, including gammaretroviruses, Achimota pararubulavirus, Quezon virus, and two adenoviruses (Baker, et al. 2013; Arai, et al. 2016; Hayward, et al. 2020). To address this, the Twist Custom Panel (Panel Name: TE-93228044_3X_FASTA; 2,256 probes; #101001) was designed with the addition of genomes (Supplementary Table 5) to the Twist Comprehensive Viral Research panel (#103545). Probes were used with the Twist Enzymatic Library Preparation Kit 1.0 (#101059), Twist Standard Hyb and Wash Kit (#101279).

#### Sequence assembly

Partial sequences include VP1, which was generated from contigs (n=2, 1,248bp and 1,959bp) with an empty string (n=60) inserted in the gap between the contigs. The partial sequence for VP4 was generated from contigs (n=3, 450bp, 315bp and

468bp) concatenated into one sequence, with two empty strings inserted into the gap between the first and second contig (n=114), and between the second and third contig (n=549).

The final partial sequence from M05058 was VP7, consisting of two contigs (n=2, 264bp and 244bp) merged into one sequence with an empty string (n=257) inserted into the gap between the two contigs.
